# Non-cell-adhesive hydrogel promotes formation of human blastoids from primed human pluripotent stem cells

**DOI:** 10.1101/2022.06.23.497328

**Authors:** Satoshi Imamura, Xiaopeng Wen, Shiho Terada, Akihisa Yamamoto, Kaori Mutsuda-Zapater, Kyoko Sawada, Koki Yoshimoto, Motomu Tanaka, Ken-ichiro Kamei

## Abstract

Artificial human blastoids are used to investigate early embryo development, pregnancy failures, and birth deficiencies, previously impossible without human embryos. Recent methods generating blastoids used human naive pluripotent stem cells, which are prone to genomic instability during *in vitro* culturing. Here, we introduce a simple, robust, and scalable method for generating human blastoids from more stable, primed human pluripotent stem cells (hPSC). Using a non-cell-adhesive hydrogel, hPSC aggregates formed an asymmetric blastoid structure with a cellular distribution similar to that of a human blastocyst, with *in vitro* implantation capability. Single-cell RNA-seq followed by cellular trajectory analysis revealed that hydrogel promoted differentiation to tri-lineage cells associated with a blastocyst. This model will allow studies on the underlying mechanisms of human pre- and postimplantation processes, consider elaborating on the potential implications of the model on assistive reproductive technology.

**One Sentence Summary:** No more than 125 characters and spaces.

---

A blastocyst is a cell aggregate with a cyst developed from the fertilized egg of a mammal, approximately 5 to 8 days after fertilization. It consists of epiblast (EPI), trophectoderm (TE), and primitive endoderm (PE) lineages. After blastocyst implantation, the TE and PE lineages give rise to the placenta and yolk sac, respectively (*1*). Conversely, the EPI lineage will form all three germ layers (e.g., ectoderm, mesoderm, and endoderm) that differentiate most types of tissue cells in the body (*2*), germ cells, and embryonic stem cells (ESCs). This structure, with its unique cell positioning, is essential for further embryonic development, tissue-specific lineage cell differentiation, and organ as well as body structure formation. The potential uses of blastocysts are four-fold. They can 1) aid the study of the early cell development, 2) be used in drug discovery for pregnancy failure (e.g., repeated implantation failure (*3*)), 3) be used to prevent birth deficiencies (*4*), and 4) be used for the advancement of assisted reproduction technology (*5*).

Unfortunately, it is difficult to obtain blastocysts from fertilized eggs for fundamental research. Moreover, using blastocysts at the industrial level is hampered by limited cell sources, donors, and ethical concerns. However, pluripotent stem cells, have great potential for applications in regenerative medicine and drug discovery. These cells include ESCs (*2*) and induced pluripotent stem cells (iPSCs) (*6, 7*). They also serve as a cell source for studying the pluripotency of stem cells *in vitro* (*8, 9*). However, the other processes of human embryonic development (e.g., blastocyst development and the implantation process) are largely unknown, compared to those of experimental animals (e.g., mice); this is due to a lack of *in vitro* research tools, limited *in vivo* access, and ethical issues. Contemporary studies have reported methods involving the use of “blastoids” where blastocyst-like structures were generated in mouse models (*10*–*12*). Moreover, recently, three groups reported the generation of human blastoids (*13*–*16*). Liu et al. used cells during the reprogramming process from fibroblasts to human pluripotent stem cells (hPSCs), as these cells contained a variety of cells, including cells with EPI-, TE-, and PE-like transcriptional signatures (*13*). Yu et al. (*14*) and Yanagida et al. (*15*), however, used naive hPSCs to generate blastoids, since recent studies have shown that naive hPSCs can differentiate into EPI-, TE-, and PE-like cells. However, naive hPSCs show genomic instability during culturing because of the hypomethylated state of the genome (*17, 18*). The culturing conditions for naive hPSCs are still being optimized, compared to those of primed hPSCs. Since the most commonly used hPSCs show primed states, it would be beneficial to use primed hPSCs to directly generate blastoids.

To generate blastoids, we focused on the physical environment surrounding natural human blastocysts. We found that human blastocysts were surrounded by a zona pellucida (ZP) glycoprotein layer, which provides a unique physical environment with 3.39 ± 1.86 kPa of Young’s modulus (*19*). Moreover, it has been shown that primed hPSCs convert to the naive-like state when three-dimensional (3D) culturing is applied (*20*). Therefore, we used a hydrogel (HG) that allows tuning of the extracellular environmental properties to mimic the ZP layer and convert the primed hPSCs to the naive state, leading to blastoid generation. Moreover, we also hypothesized that the asymmetric structure of blastocysts could be generated by local concentration gradients of growth factors and cytokines; thus, the HG would be suitable for generating such gradients.

Here, we developed a method to generate blastoids, specifically HG blastoids (HG-blastoids), from primed hPSCs in an HG medium through 3D cell culturing. This method uses a thermoresponsive HG (a copolymer of poly(*N*-isopropyl acrylamide) and poly(ethylene glycol) (PNIPAAm-PEG)) (*21, 22*), which holds hPSC aggregates with the stimulation of elastic force against the growth thereof, without adhering to the cells. Additionally, because it can perform a sol-gel transition *via* temperature change, the HG allows the mixing and harvesting of cell aggregates from the solution at a low temperature (<20 °C). Furthermore, 3D cell culturing in the HG-containing medium can be performed at 37 °C. Since serum contains a variety of growth factors for inducing differentiation of hPSCs to trophoblasts (*23*) and maintaining hPSC selfrenewal, it would allow for the generation of asymmetric blastocyst-like cysts. After generating HG-blastoids, we conducted a variety of assays to determine their cellular identities and locations within the HG-blastoids as well as investigate their functionalities, such as derivation of pluripotent/trophoblast stem cells and implantation, as an *in vitro* model of a human blastocyst.

## Generating human blastoids in hydrogel

To generate human HG-blastoids from primed hPSCs, we created a physical environment using a HG, to mimic a ZP glycoprotein layer (**Fig. 1A**). We used KhES1 hESCs expressing an EPI marker fused to the enhanced green fluorescent protein (eGFP), downstream of the octamer-binding transcription factor 4 promoter (OCT4; or POU domain, class 5, transcription factor 1 [POU5F1]), to visualize cells expressing OCT4 and ICM-like cell aggregates in blastoids. The EPI marker used was called K1-OCT4-eGFP (*24*). We used mTeSR-1 medium to maintain primed hPSCs (*25, 26*). In terms of the HG, we found that 10 % (w/v) of PNIPAAm-PEG (HG), which provides 0.15 ± 0.04 kPa of Young’s modulus, allows culturing of primed hPSCs (*21*) (**fig. S1 and table S1**); hence, we used this HG for blastoid generation.

**Fig. 1.**
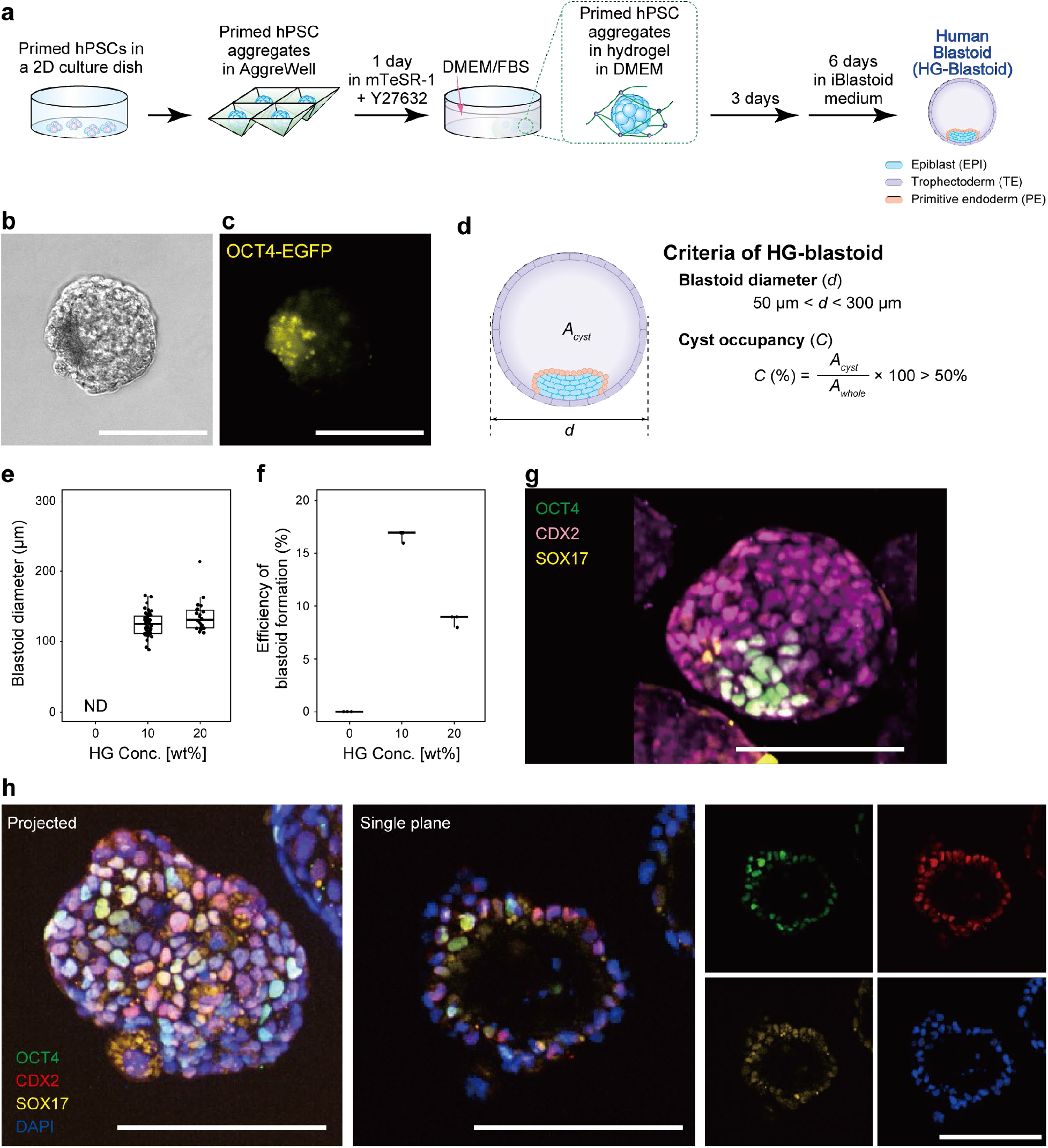
Principle for generating blastoids from human primed pluripotent stem cells (hPSCs). **(A)** A schematic showing the experimental process for generating HG-blastoids from primed hPSCs. TB, trophoblasts; ICM, inner cell mass; PE, primitive endoderm. **(B)** Phase-contrast and **(C)** fluorescent micrographs of an HG-blastoid derived from K1-OCT4-eGFP cells. **(D)** Illustration of the criteria to identify HG-blastoids among the other hPSC aggregates. “*d*” shows the diameter of HG-blastoids. “*A*_*cyst*_” and “*A*_*whole*_” indicate the area of cyst and whole aggregate, respectively. **(E)** Size distribution of HG-blastoids in 0, 10, and 20%(w/v) HG at the end of HG-blastoid generation process shown in **Fig. 1A**. Center lines show the medians; box limits indicate the 25^th^ and 75^th^ percentiles; whiskers extend 1.5 times the interquartile range from the 25^th^ and 75^th^ percentiles. **(F)** Generation efficiency of HG-blastoids in 0, 10, and 20%(v/v) HG. The lines show the minimum, median and maximum values. **(G, H)** Whole-mount immunocytochemistry micrographs for the observation of typical markers of epiblast (OCT4), trophectoderm (CDX2), and primitive endoderm (SOX17) in HG-blastoids. DAPI was used to stain nuclei in image **Fig. 1E**. Scale bars represent 100 μm.

To obtain hPSC aggregates of uniform size (and to increase reproducibility), single-cell-dissociated primed K1-OCT4-eGFP hESCs cultured in a 2D-culture dish were transferred to an AggreWell plate (*27, 28*) and cultured in mTeSR-1 medium supplemented with 10 μM Y27632 ROCK inhibitor (*29*) for 1 day. Next, K1-OCT4-eGFP cell aggregates were harvested from the AggreWell plate, placed in DMEM supplemented with 10 or 20 % (w/v) HG solution at 4 °C, and transferred to an ultralow attachment cell culture dish at 37 °C. Next, the hESC aggregates encapsulated in HG were cultured in DMEM supplemented with 10 % (v/v) FBS. After 3 days of culturing, the culture medium was replaced with iBlastoid medium (*13*) and cultured for a further 6 days to form HG-blastoids. Both 10 and 20% HG conditions provided blastoids with a small portion of cell aggregates expressing the *OCT4* promoter-driven eGFP, representing the ICM-like structure (**Fig. 1B, C and fig. S2a**). We established as inclusion criteria those HG-blastoids with a diameter ranging between 50–300 μm with a cyst occupancy of over 50% (**Fig. 1D**).

While both 10 and 20% HG conditions generated the blastoids with similar diameters (124.2 ± 16.5 and 134.4 ± 20.9 μm, respectively; **Fig. 1E**), the 10% (w/v) HG conditions showed a 16.6% higher efficiency to generate HG-blastoids (60 out of 300 hPSC aggregates) than that of 20% (w/v) HG (8.6 ± 0.5%; **Fig. 1F**). By using this 10% (w/v) HG condition, we examined the efficiency to generate blastoids of each initial seeding cell numbers in a well of the AggreWell 400 plate. We identified 1.5 × 10^5^ cells as the optimal cell number (**fig. S2b and table S2**), while no significant differences were observed in the blastoid sizes among the tested initial seeding cell numbers (**fig. S2c**). We also tested other hydrogels with similar properties, such as agarose (**fig. S2d-f, and table S2**), which is non-adherent to cells and has a tunable stiffness (**fig. S1**). We confirmed that agarose also had the capability to support the generation of HG-blastoids, although the efficiency was much lower than that of PNIPAAm-PEG [7.7 ± 4.8% with the use of 0.5%(w/v) agarose; **fig. S2f**]. Interestingly, when using collagen gel (2 mg/ml; 0.15 kPa of Young’s Modulus(*30*)), the aggregates were expanded into the gels without maintaining their structure 24 hours after introduction (**fig. S2g**).

To confirm the cellular distribution representing a human blastocyst in HG-blastoids, we observed the expression patterns of typical markers, including OCT4 (epiblast), CDX2 (trophectoderm), and SOX17 (primitive endoderm) (**Fig. 1G, H, fig. S3a, and table S3**). These results indicated that the obtained HG-blastoids mimicked the cellular distribution of the three lineages in the natural blastocyst. In addition, HG conditions with serum induced the differentiation of cells at the outer layer of cell aggregates, whereas the cells within the aggregates maintained *OCT4* expression.

In addition to HG-blastoids derived from primed K1-OCT4-eGFP hESCs, we also tested primed 585A1 hiPSCs(*31*) for HG-blastoid formation. By using the 10% HG condition and 1.5 × 10^5^ cells of the initial cell seeding density, 585A1 cells were able to form HG-blastoids with 131.5 ± 8.9 μm in a diameter, and showed 23.11 ± 10.70% formation efficiency (**fig. S2h**). The obtained HG-blastoids from hiPSCs had the expression patterns of typical markers, including OCT4, CDX2, and SOX17 (**fig. S3b**)

### Cells in HG-blastoids showed similar gene expression signatures and cellular populations as the natural human blastocyst

To investigate cellular identities in HG-blastoids, single-cell RNA sequencing (scRNA-seq) (*32, 33*) was performed using approximately 400 HG-blastoids derived from K1-OCT4-eGFP hESCs containing 5,559 cells (**Fig. 2A**). We showed that the HG-blastoids had three lineages, as identified by their corresponding gene markers (*POU5F1, NANOG, SOX2*, and *NODAL* for EPI cells; *GATA2, GATA3*, and *ABCG2* for TE cells; *GATA6, FN1*, and *COL4A1* for PE cells; **Fig. 2B, C and fig. S4**). Although hESCs have two major cell populations, they strongly express *POU5F1*. Compared with hESC transcriptomic signatures, a cluster shown in light blue in **Fig. 2A** was observed only in HG-blastoids, unlike hESCs, which showed TE- and PE-like transcriptomic patterns. Similar to **Fig. 2A**, we also found some extracellular populations, which could not be assigned to EPI-, TE-, or PE-like cells, and thus were named “Intermediated (IM)” cells. We also compared the scRNA-seq data with that of the original hESCs (13,516 cells), as well as 585A1 hiPSCs (1,423 cells) and their HG-Blastoids (998 cells) (**fig. S5**). We observed an overlap of some clusters between original hESCs/hiPSCs and HG-blastoids, but HG-Blastoids had some distinguishable cell populations, which showed TE- and PE-like gene expression signatures. Importantly, generated HG-Blastoids from both hESCs and hiPSCs showed similar cellular populations (**fig. S5**).

**Fig. 2.**
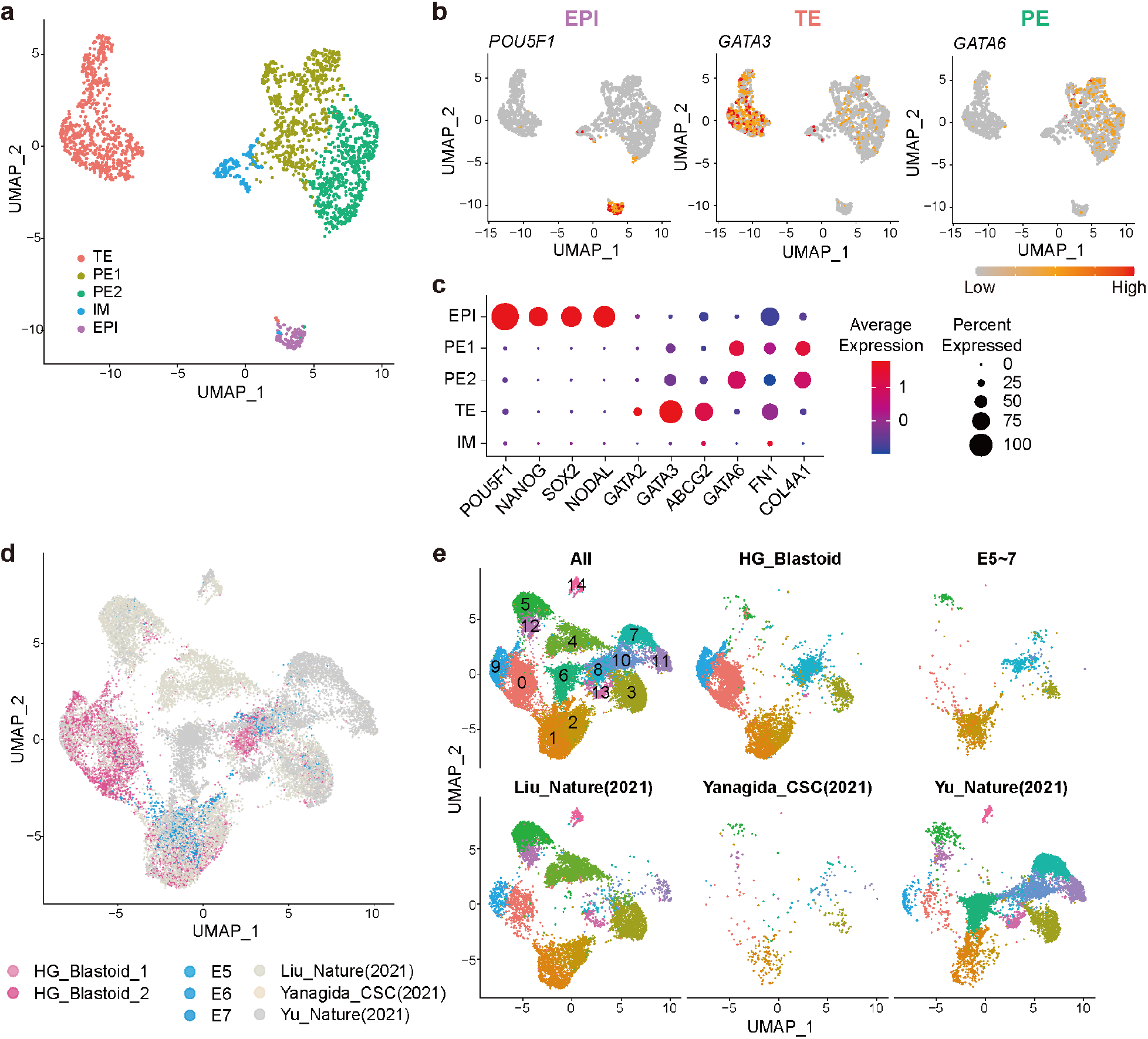
Single-cell transcriptomic analyses of HG-blastoids. **(A)** UMAPs of the 2,072 cells in the sc-RNA-seq library of HG-blastoid derived from K1-OCT4-eGFP hESCs. **(B)** Expression of EPI (*POU5F1*), TE (*GATA3*), and PE (*GATA6*) marker genes. **(C)** A dot plot showing the expression of marker genes for EPI (*POU5F1, NANOG, SOX2*, and *NODAL*), TE (*GATA2, GATA3*, and *ABCG2*), and PE (*GATA6, FN1*, and *COL4A1*). Dot size and color (blue to red) represent the percentage of cells with the gene expression and averaged expression values. **(D, E)** Integrated and split UMAPs to show integrated scRNA-seq data sets consisting of data from HG-blastoids, human E5-E7 blastocysts (*34*), and other published studies on blastoids (*13*–*15*).

Next, we used comparative transcriptomics to determine whether the EPI-, TE-, and PE-like cells in HG-blastoids derived from K1-OCT4-eGFP hESCs showed similarities in transcription signatures to natural human blastocysts. Here, scRNA-seq data of human blastocysts on days 5–7 (E5-7) ((*34*)) was compared to our data (**Fig. 2D**, and **fig. S6**). Considering the integrated UMAP, HG-blastoids had EPI-(Clusters #3, #8, and #10), TE-(Clusters #1 and #2), and PE-like cells (Cluster #5) assigned to E5-7 cellular populations by their transcriptomic signatures. Next, an enrichment analysis of gene ontology (GO) terms was performed for each cluster (**figs. S7** and **S8**). EPI-like clusters had “signaling pathways regulating pluripotency of stem cells (ko04550),” “embryonic morphogenesis (GO:0048598),” and “cell division (GO:0051301),” which are important functions of EPI. In TE-like clusters, the GO term of “placenta development (GO:0001890)” was significantly enriched, indicating the potential of TE differentiation to the placenta, which is a critical process of blastocyst implantation. PE-like clusters showed an enrichment in “endoderm formation (GO:0001706),” indicating PE formation.

Although a similar percentage of cells was observed in both HG-blastoids and E5-7 blastocysts, HG-blastoids showed fewer populations of TE- and PE-like cells (**fig. S9**). More specifically, E5-7 blastocysts had 24.2 %, 65.2 %, and 20.5 %, of EPI-, TE-, and PE-like cellular populations, respectively, while HG-blastoids had 21.4 %, 27. 9%, and 1.08 %. However, HG-blastoids had additional cellular populations. For example, cells in cluster #0 were present in both human blastocysts and HG-blastoids, although to a lesser degree in human blastocysts, 4.69 % vs. 34.0 % in HG-blastoids (**fig. S9**). According to the GO enrichment analysis, “appendage development (GO:0048736)” and “cell division (GO:0051301)” were highlighted in HG Blaastoids. With regard to cluster #9, the GO term “Extracellular matrix organization (R-HAS-1474244)” was enriched. Because HG physical stimuli were introduced to generate HG-blastoids, cells might be involved in the process of ECM (extracellular matrix) remodeling. These results suggest two possible future directions. One is to improve the efficacy of direct differentiation of the TE and PE lineages with both chemical and physical stimuli, while the second is to collect blastoids. It is important to note that we collected HG-blastoids by evaluating their microscopic morphologies, without analyzing their molecular signatures. This may cause contamination with other cell aggregates with partial induction toward blastoid generation, resulting in single-cell profiles.

Thus, the development of HG-blastoids similar to natural human blastocysts needs further improvement.

Finally, we compared the transcriptomic signatures of individual cells in HG-blastoids with those published by the other groups in 2021. These groups reported the generation of blastoids *in vitro* (*13*–*15*). By integrating our scRNA-seq data sets with theirs, along with data from human E5–E7 blastocysts (*34*), we investigated the differences and advantages over the other methods (**Fig. 2D, E**). UMAP integration of these datasets revealed that all methods provided blastoids with EPI-, TE-, and PE-like clusters; however, the cellular populations were quite different (**fig. S9**).

Although the number of cells (306) in the dataset of Yanagida et al. (2021) was much smaller than the rest (7,778 cells (*13*); 6,840 cells (*15*); our data [5,559 cells]), Yanagida’s blastoids had cells assigned to EPI, TE, and PE clusters. Moreover, their blastoids showed close similarities to the cellular profiles of human E5-7 blastocysts. Conversely, blastoids generated by Liu et al. (*13*) and Yu et al. (*15*) showed more heterogeneous populations, wherein E5-7 blastocysts were absent. Moreover, a key difference was found in cluster #11. This cluster was absent in HG-blastoids and E5-E7 blastocysts but present in the other blastoids, ranging from 1 % to 9 % of cells. Furthermore, cluster #11 showed apoptotic cells with enriched GO terms related to apoptosis (e.g., “apoptotic signaling pathway [GO:0097190]” and “positive regulation of cell death [GO:0010942]”), suggesting that their method may cause cellular damage during blastoid formation. Our method, however, with the use of HG, provides more suitable conditions without harmful cell damage.

### HG maintains differentiation into HG-blastoid lineage

To investigate the mechanisms of how HG contributes to blastoid generation from primed hESCs, we carried out scRNA-seq followed by pseudotime analysis (*35, 36*) for HG-blastoids derived from K1-OCT4-EGFP hESCs on days 0, 4, 6, and 9 as well as the hESC aggregates without HGs on days 4 and 6 (**Fig. 3**). This dataset showed 11 clusters, including four EPI (EPI1–4), two TE (TE1 and 2), two PE (PE1 and 2), and three Mixed clusters (Mix1, 2 and 3) (**Fig. 3A and table S5**). Among EPI clusters, cells in EPI1 expressed the EPI markers (i.e., *OCT4, SOX2, NANOG*, and *NODAL*) more than the other EPI clusters (**fig. S10a**). In the case of TE clusters, although *GATA3* and *KRT7* in both TE clusters were highly expressed, *GATA2* and *CDX2* were expressed at the same levels as observed in EPI, PE, and Mix clusters (**fig. S10b**). In PE clusters, *GATA6, COL4A1*, and *PITX2* were highly expressed in both PE clusters, but *GATA4* was not expressed in all clusters. (**fig. S10c**). Besides EPI, TE, and PE clusters, we found three clusters, which could not be categorized cells by the expression of EPI, TE, and PE markers. In light of pseudotime analysis, pseudotime around 10 could be considered at the initial time point to start blastoid generation, since hESCs on day 0 had EPI1, EPI2, EPI3, EPI4, Mix1, and Mix2. Then, PE1 and TE2 appeared followed by TE1 and Mix3 (**Fig. 3B-D**).

**Fig. 3.**
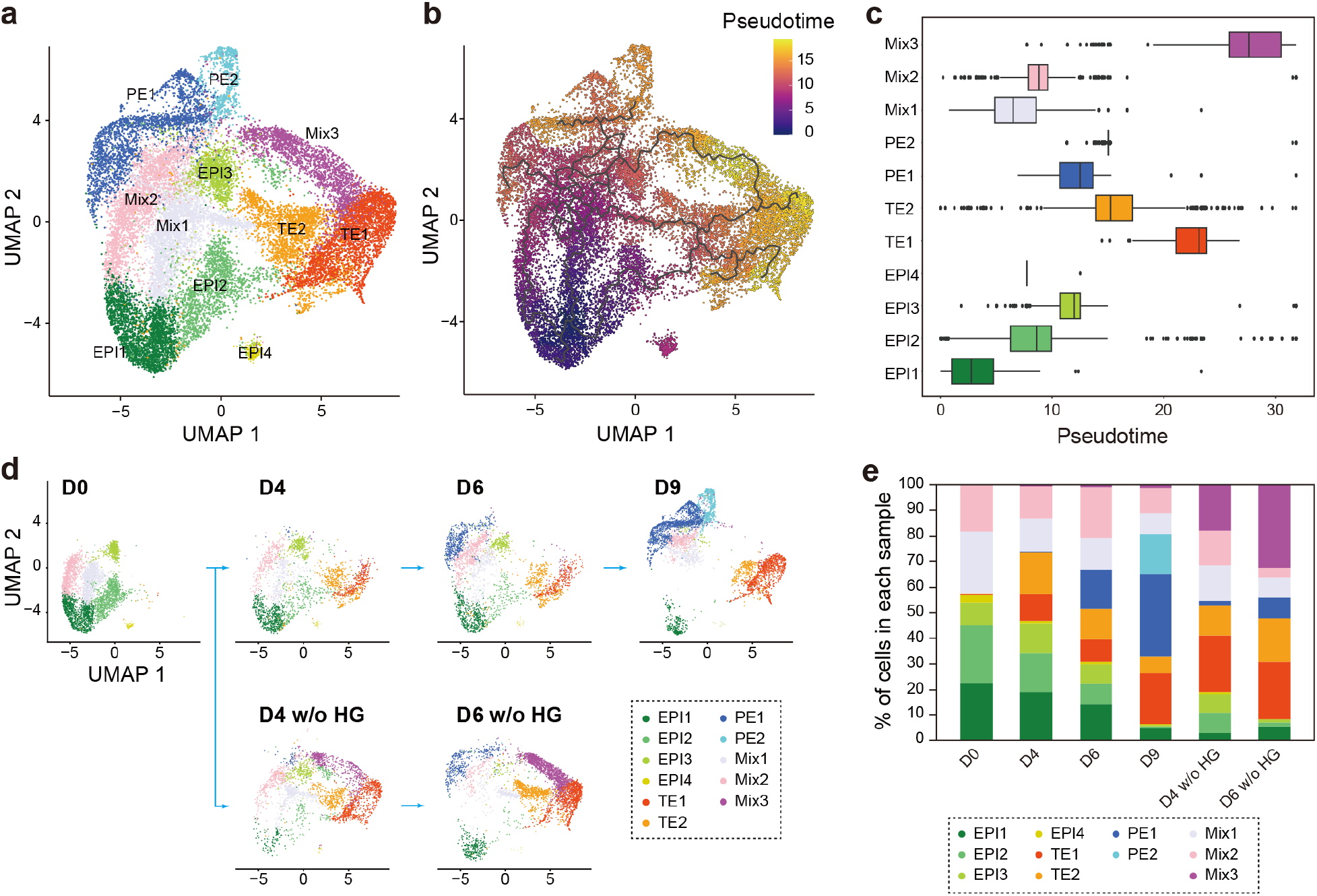
Single-cell trajectory analysis to investigate the role of HG during blastoid generation from primed hESCs. **(a)** Integrated UMAP of K1-OCT4-EGFP hESC aggregates treated with HG on days 0, 4, 6, and 9. As control, K1-OCT4-EGFP hESC aggregates without HG (D4 w/o HG and D6 w/o HG) were also used for integrated scRNA-seq analysis. **(b)** UMAP projected with pseudotime. **(C)** Box plot to show the cellular distribution of each cluster along pseudotime. Center lines show the medians; box limits indicate the 25^th^ and 75^th^ percentiles; whiskers extend 1.5 times the interquartile range from the 25^th^ and 75^th^ percentiles. **(D)**, Split UMAPs of K1-OCT4-EGFP hESC aggregates treated with HG on days 0, 4, 6, and 9 as well as K1-OCT4-EGFP hESC aggregates without HG (D4 w/o HG and D6 w/o HG). **e**, Percentiles of clustered cells in each sample.

We used HG to generate the blastoids from primed hPSCs. Primed hESCs on day 0 had 57.0% of cells associated with EPI clusters (EPI1, EPI2, EPI3 and EPI4), and 42.6% of cells with the mixed cellular populations (Mix1 and 2) (**Fig. 3D, E**). During the blastoid generation process with the use of HG, EPI2, EPI3, and EPI4 clusters gradually disappeared, and only the EPI1 cluster remained at 5.0% of total cellular population on day 9. In contrast, TE clusters were not observed on day 0, but the TE1 cluster was 10.3% of cellular population on day 4, and increased to 20.0% on day 9, while the TE2 cluster rapidly appeared 16.4%, and then reduced to 6.4% on day 9. In the case of PE lineage, cells associated with PE clusters appeared around 0.5% on day 4, and increased to 15.4% and 47.9% on days 6 and 9, respectively. Until day 6, only cells in PE1 appeared, and cells in the PE2 cluster were observed on day 9.

For comparison, we also investigated the changes of cellular populations within hESC aggregates without the use of HG. In this condition, hESC aggregates did not form a blastocyst-like structure (**fig. S2**). Therefore, we harvested hESC aggregates without any selection by morphologies of cell aggregates for scRNA-seq. Similar to HG-blastoid generation, hESCs without HG on days 4 and 6 showed the dramatic reduction of cell numbers in EPI1 (2.9% and 5.5%), EPI2 (7.9% and 1.5%) and EPI4 (0.9% and 0.1%) as hESC with HG. Cells in EPI3 showed the reduction of cell numbers on day 6 (1.4%), while cell numbers did not change drastically until day 4. The most distinguishable cluster was Mix3. This cluster notably appeared in only hESC aggregates without HG on days 4 and 6, but not under any other conditions (**Fig. 3D, E**). Therefore, we extracted genes highly expressed in Mix3 and used CellMaker 2.0 to more thoroughly investigate candidate cell types (**table S6**). The results showed that Mix3 expresses various cell markers, including markers related to neurons and astrocytes that constitute the brain, astrocytes that constitute the embryo body, and gonadal endothelial cells, granulosa cells, and mitotic fetal germ cells that constitute the gonads.

### HG-blastoids are capable of providing naive/primed PSCs and TSCs

Natural blastocysts provide pluripotent and trophoblast stem cells (PSCs (*2*) and TSCs (*37*), respectively). To evaluate whether the generated HG-blastoids could produce these stem cells, we digested the HG-blastoids and cultivated the cells under the culture conditions for naive/primed PSCs and TSCs, namely naive/primed bPSCs and bTSCs, respectively (**Fig. 4A**). The obtained naive bPSCs showed a typical semi-spherical colony morphology and expressed KLF4, NANOG, and OCT4 (*38*) (**Fig. 4A, fig. S11a, and table S3**). The obtained primed bPSCs showed a flat colony morphology and expressed TRA-1-60, OCT4, and NANOG (**Fig. 4C, fig. S11b, and table S3**). The original hESCs used in this study were maintained in a primed PSC culture medium, mTeSR-1 (*26*); however, the obtained HG-blastoids showed the capability of providing all three stem cells, suggesting that our method to generate blastoids is also a new method to obtain naive bPSCs from primed hPSCs. While maintaining TSCs (*37*), bTSCs expressed KRT7, CDX2, and GATA2, markers associated with TSCs (*37*) (**Fig. 4D, fig. S11c, and table 3**). Thus, we conclude that the HG-blastoid can produce the same set of stem cell lineages as a natural human blastocyst.

**Fig. 4.**
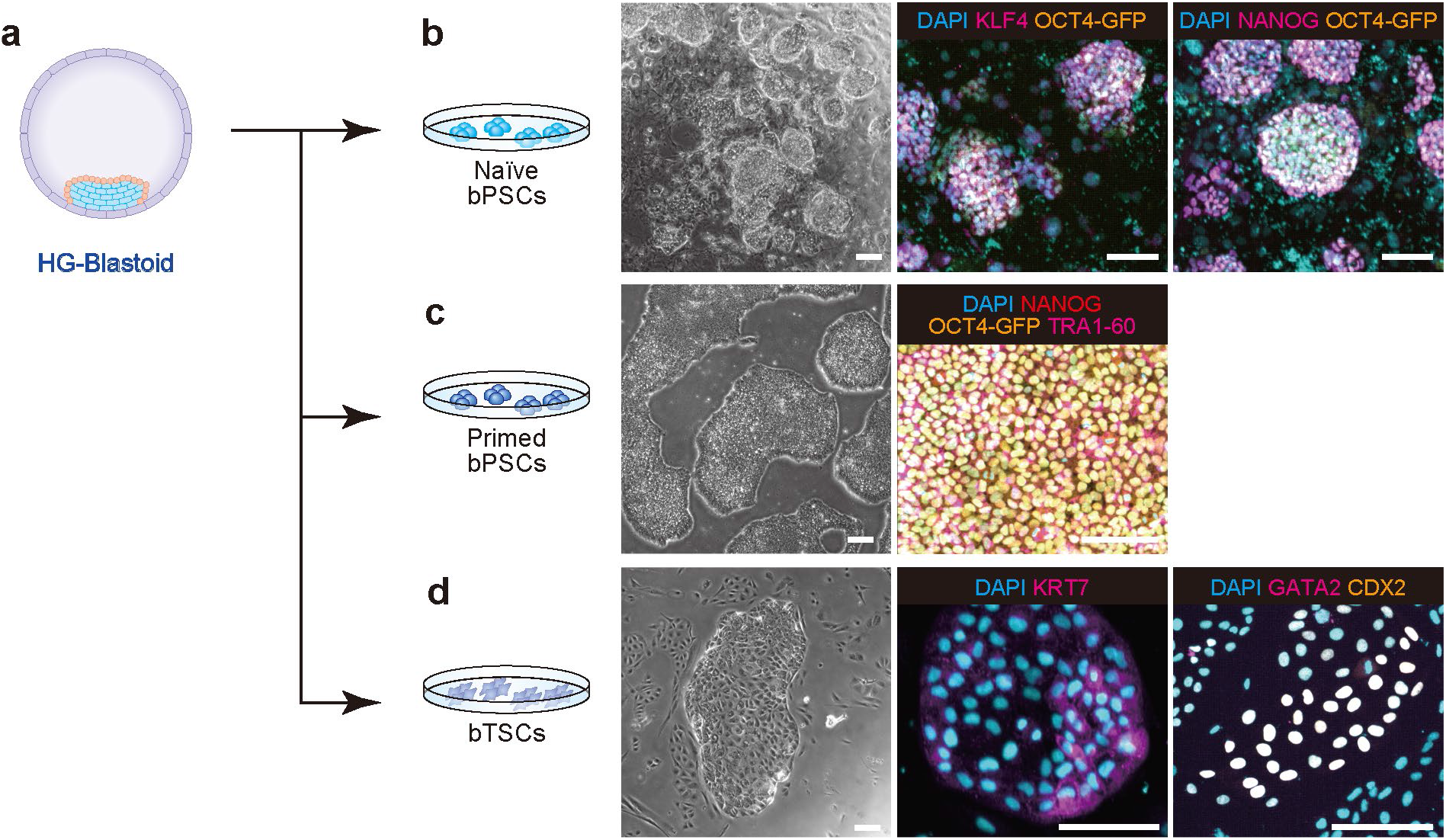
Derivation of stem cells from HG-blastoids. **(A)** Blastocysts provide three kinds of stem cells, including naive/primed pluripotent stem cells (bPSCs) and trophoblast stem cells (bTSCs). Phase-contrast and fluorescent micrographs of immunocytochemistry of **(B)** naive bPSC visualized using KLF4, NANOG, and OCT4-eGFP, **(C)** primed bPSCs visualized using NANOG, TRA1-60, and OCT4-Egfp, and **(d)** bTSCs visualized by GATA2 and CDX2. DAPI was used to stain nuclei. Scale bars represent 100 μm.

### HG-blastoids show the implantation features *in vitro*

During the human implantation process, trophoblasts of a blastocyst attach to the uterine wall, and 4 days after adhesion, the inner cell mass turns to the pro-amniotic cavity and epiblasts. To evaluate whether human HG-blastoids showed such blastocyst functionality, HG-blastoids were transferred onto the cell culture dish coated with GelTrex extracellular matrices to recapitulate the implantation process *in vitro* (**Fig. 5A**). After attachment, the HG-blastoids gradually expanded and grew on the substrate (**Fig. 5B**).

**Fig. 5.**
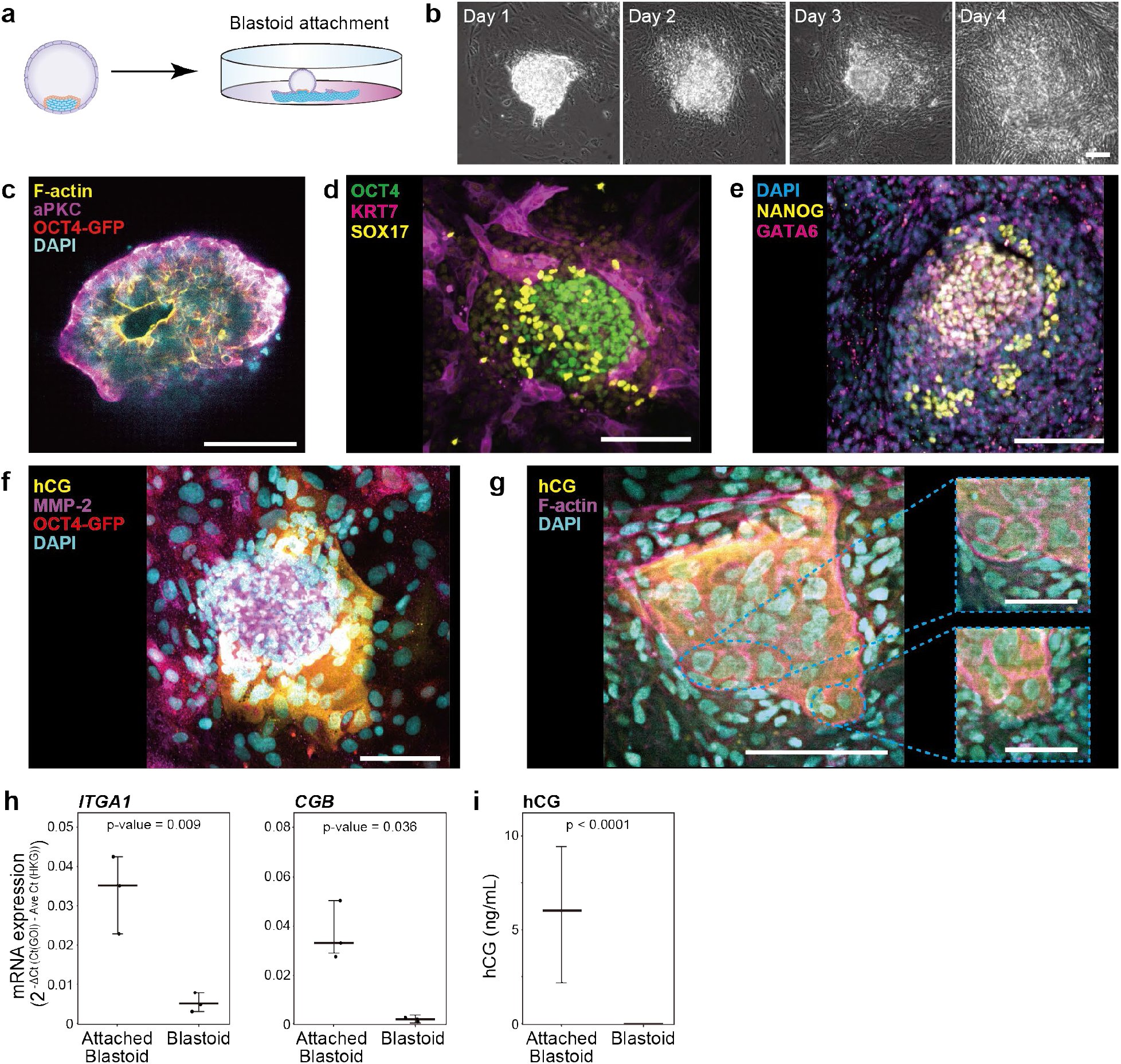
*In vitro* implantation assay for HG-blastoids. **(A)** A schematic showing the *in vitro* implantation assay, whereby HG-blastoids were seeded onto the GelTrex-coated cell culture dish. **(B)** Micrographs of HG-blastoids attached to the dish. **c-f**, Fluorescent micrographs of attached HG-blastoids, 4 days after *in vitro* implantation, immunostained with the following: **(C)** F-actin, OCT4, and aPKC; **(D)** OCT4, SOX17, and KRT7, to investigate the distribution of EPI-, PE- and TE-like cells, respectively; **(E)** NANOG and GATA6; **(F)** Hcg and MMP2 (markers of extravillous cytotrophoblasts [EVTs] and syncytiotrophoblasts [STs], respectively); **g**, Hcg and F-actin. OCT4-GFP represents Egfp expression driven by the *OCT4* promoter. DAPI was used to stain nuclei in images **(C, E, F**, and **G)**. Scale bars represent 100 μm, except scale bars in images **g** with dashed light blue squares represent 50 μm. **(H)** Quantitative RT-PCR analysis of gene expression associated with EVTs (e.g., *ITGA1*) and STs (*CGB*; also known as hCG). n = 3. **i**, Quantification of hCG secreted from HG-blastoids and attached HG-blastoids. n = 3.

In light of previous reports, an *in vitro* implantation assay for a human blastocyst is capable of recapitulating the formation of a lumen-like pro-amniotic cavity (*39, 40*). The attached HG-blastoids formed a pro-amniotic-like cavity, in which F-actin and aPKC were expressed in the lumen region, while OCT4-positive EPI-like cells were expressed around the lumen, as confirmed by immunocytochemistry (**Fig. 5C and table S3**). Moreover, co-staining experiments using OCT4 (EPI), KRT7 (TE), and SOX17 (PE), revealed that OCT4-positive cells showed cell aggregates with high cellular densities, a feature of EPI. Furthermore, SOX17-positive PE-like cells were located on the side of EPI-like cell aggregates, while KRT7-positive TE-like cells showed outgrowths from the attached areas (**Fig. 5D and table S3**). As shown in **Fig. 5E**, the cells expressing the GATA6 PE marker were next to the NANOG-positive EPI-like cell aggregates, a feature of PE-like cells. These results were in agreement with the staining results shown in **Fig. 5D**.

To evaluate TE-like features in detail, MMP2 and hCG were stained as representative markers of extravillous cytotrophoblasts (EVTs) and syncytiotrophoblasts (STs), respectively (**Fig. 5F, G and table S3**). Close observation of hCG-positive ST-like cells and their nuclei, showed some cells with multiple nuclei, representing a unique feature of ST-like cells (*41*) (**Fig. 5G**).

Conversely, MMP2 positive cells showed spindle-like morphologies, a cellular phenotype of EVT-like cells (*41*). Next, RT-PCR was used to quantify the expression of genes associated with EVT- and ST-like cells. All of the EVT marker genes (e.g., *ITGA1, ITGA5*, and *FN1*) were highly induced after attachment, while only the *hCG* ST-associated genes (but not *CSH1* and *SDC1*) were induced (**Fig. 5H, fig. S12, and table S4**). These results suggested that ST-like cells generated from HG-blastoids were partially functionalized as STs. Moreover, the secretion of hCG (a chemical secreted from trophoblast tissues, detected during normal pregnancy) was detected in the culture medium 4 days after *in vitro* implantation of HG-blastoids (**Fig. 5I**), suggesting that trophoblast-like cells appeared in attached HG-blastoids as those of natural human blastocysts.

In 2021, the International Society for Stem Cell Research (ISSCR) updated its guidelines for stem cell research and clinical translation (*42*). The ISSCR recommends that research with *in vitro* embryo models containing the epiblast and extraembryonic endoderm, covered by trophectoderm-like cells (e.g., blastoid), would not be permitted in *in vitro* culture beyond the appearance of the primitive streak. Therefore, we measured the expression of *TBXT, EOMES*, or *MIXL1* as primitive streak markers (*43, 44*) in the attached HG-blastoids in a dish for 4 days, and we confirmed that HG-blastoids did not express these markers, as confirmed by quantitative RT-PCR (**fig. S13 and table 4**).

## Discussion

We established a simple method to generate blastoids from human primed PSCs by applying the proper physical environment with hydrogels (HG) to mimic that of natural human blastocysts. The obtained HG-blastoids showed similar patterns of cellular distribution (**Fig. 1**), molecular expression signatures (**Fig. 2**), and the capability to provide naive/primed PSCs and TSCs (**Fig. 4**), as natural human blastocysts. Moreover, they demonstrated implantation capability *in vitro* (**Fig. 5**). These capabilities illustrate its potential as a powerful tool for studying human early embryonic developmental processes *in vitro* (*16, 45*).

To generate human blastoids from primed hESCs, we used a HG capable of mimicking the physical extracellular environment. A natural human blastocyst has a ZP glycoprotein layer, which functions as the selection layer for sperm and immunocontraception. The biochemical and physical properties of the ZP layer are well known (*46*–*50*). Particularly, it is known that the physical properties of the ZP layer change after fertilization, with zona hardening critical for preventing polyspermic penetration (*47*). The HG in this study showed 0.15 ± 0.04 kPa of Young’s modulus (**fig. S1 and table S1**), which is softer than the reported Young’s modulus of a natural ZP layer (ca. 10 -40 kPa) (*19, 48*–*50*). When we applied 2.01 ± 0.10 kPa for blastoid formation of primed hESCs, no blastoids were obtained. Without the HG, hESC aggregates in the cell culture medium with 10 % (v/v) serum did not form blastocyst-like structures but rather formed embryoid bodies (EB) that contained three germ layers (ectoderm, mesoderm, and endoderm). This method is traditionally used to induce the differentiation of hPSCs into the desired tissue cells; however, the generated EBs have random cell distributions. When we applied the HG with 10 % (v/v) serum, the outer layer of the hPSC aggregates received the biochemical and physical stimuli, inducing differentiation to the trophectoderm lineage. The cells inside the hPSC aggregates, however, did not receive these stimuli and maintained their EPI phenotype. Therefore, although the HG used for blastoid formation was softer than that of the ZP layer, such a physical environment is critical for obtaining blastoids from primed hPSCs. Since the harder ZP layer mainly prevents massive sperm penetration into an oocyte, it might not be necessary to generate blastoid formation. Such a layer could not adhere to cells in a blastocyst or blastoid because they could not maintain a blastocyst-like structure due to their migration.

Indeed, agarose gel also showed the capability to generate blastoids from primed hPSCs (**fig. S2d-f**). In contrast, extracellular matrix proteins (e.g., collagen) or cell-adhesive polymers (e.g., poly-L-lysine) would not be suitable to generate blastoids, as shown in **fig. S2g**. Conversely, non-adhesive polymers (e.g., agarose) support the formation of blastoids from primed hESCs. We also compared aggregate not embedded in HG with aggregate embedded in HG to clarify the direct effect of HG on blastoid formation. In light of **Fig. 3**, the Mix3 cluster appeared only in aggregates without HG, and the Mix3 cluster contained a variety of tissue cells. Since the suspension culture of hPSC aggregates in serum-contained medium shows EB, the Mix3 cluster could represent the mixture of all three germ layers toward EB formation. These results suggest that HG inhibited cell differentiation, while simultaneously inducing differentiation into cells of TE and PE and remaining EPI.

In 2021, the number of studies on human blastoid generation increased, yet the cell sources in these studies were quite diverse. Liu et al. (2021) showed that cells treated for reprogramming from fibroblasts to hiPSCs have a variety of cell types (e.g., EPI-, TE-, and PE-like cells), and have the ability to form blastoids (*13*). However, cellular heterogeneity during reprogramming is unpredictable and unregulatable, as the obtained blastoids showed more cellular components, unlike the natural blastocysts at E5–E7 (**Fig. 2D, E**). Instead of using reprogrammed cells, other reports used naive hPSCs to obtain blastoids (*14, 15*). As previously reported, the naive state is considered more undifferentiated than the primed state (*51*). Human PSCs were thought to be unable to differentiate into extraembryonic cell lineages (e.g., trophoblast stem cells for forming placenta). However, recent studies have shown that naive hPSCs can differentiate into these lineages by changing their culture conditions (*23, 52, 53*). Therefore, undifferentiated naive hPSCs were used to generate blastoids. Recent studies have shown that primed hPSCs can be converted to a naive state. However, naive hPSCs cultured in 5i/L/A show genomic instability during culturing due to the hypomethylated state of the genome (*17, 18*). The culture conditions for naive hPSCs are still being optimized compared with those of primed hPSCs. To the best of our knowledge, human blastoids have not yet been generated from primed human PSCs.

Although the underlying mechanism is still unclear, the presented approach is a promising approach to study the processes and mechanisms underlying the generation of human blastoids from primed hPSCs. Moreover, it is capable of producing naive/primed PSCs and TSCs. In mouse models, a recent study demonstrated the process of primed mESCs to blastoids (*10*) which could be a useful tool for elucidating the generation of blastoids from primed human ESCs.

Finally, the HG-blastoids were capable of undergoing an implantation process *in vitro*, showing the formation of a lumen-like pro-amniotic cavity, trophoblast tissues, and secretion of hCG, without expressing primitive streak markers (*TBXT, EOMES*, or *MIXL1*) (**fig. S13**). By using such blastoid models, we can recapitulate and study the early human embryo developmental process before the appearance of the primitive streak (*42, 54*). Although the ISSCR updated the guidelines for human embryo models derived from stem cells, a discussion is needed on how we can achieve a good balance between the advancement of the fundamental understanding of human development and ethical concerns.

This method had some limitations that need to be addressed. Although we were able to generate HG-blastoids from primed hPSCs, the efficiency was still low (around 15∼20 %) compared to that of other reported methods using naive hPSCs. We hypothesized that primed hPSCs could be partially converted to a naive state by the physical environment of the HG, and then undergo blastoid formation. Therefore, the conversion would be the rate-limiting step for blastoid formation. Additionally, selection methods for proper blastoids have not yet been well established. The selection methods and quality checks for the obtained blastoids were based on microscopic imaging to observe their morphologies. In particular, the methods used for quality checking were originally established for blastocysts by *in vitro* fertilization (IVF), with no concerns regarding contamination of other types of cells. According to the scRNA-seq results, the reported blastoids, including HG-blastoids, contained impurities in the cells, which should not exist in a natural human blastocyst. Therefore, the selection methods for high-quality blastoids require microscopic imaging to observe not only blastocyst-like morphology, but also molecular markers for EPI-, TE-, and PE-like cells.

In conclusion, we established a simple and robust method for generating human HG-blastoids from primed hPSCs treated with a physically tuned HG. We envision that this method can be further improved to develop clinically relevant and chemically defined conditions to understand the basis of early human embryonic development. This method can be extended to other species to understand the fundamentals of early embryonic development by comparing multiple species without using their embryos. Finally, it can be used for high-throughput screening in drug discovery for pregnancy failure and birth defect prevention.

## Acknowledgments

We thank Kaho Takamuro and Ayumi Kikkawa for their technical assistance. We thank Dr. Kouichi Hasegawa, Dr. Dan Ohtan Wang, Dr. Tomonori Nakamura, and Prof. Michinori Saito for critical discussion. The authors thank to iCeMS Analysis Center to access the advanced microscopy and analytical instruments. The authors thank Dr. Takefumi Kondo for carrying out RNA sequencing. The WPI-iCeMS is supported by the World Premier International Research Centre Initiative (WPI), MEXT, Japan.

## Author contributions

Conceptualization: SI, XW, KK

Methodology: SI, XW, ST, AY, MT, KK

Investigation: SI, XW, ST, AY, KMZ, KS, KY, MT, KK

Visualization: SI, XW, AY, KMZ, KK

Funding acquisition: KK

Project administration: KK

Supervision: KK

Writing – original draft: SI, XW, KK

Writing – review & editing: SI, XW, KK

## Data and materials availability

All sc-RNA-seq data are available at the Gene Expression Omnibus (GSE206469, GSE231484, GSE231683).

## Supplementary Materials

Materials and Methods

figs. S1 to S13

tables S1 to S6

References (1–16)

