## Supplementary information for "Non-cell-adhesive hydrogel promotes formation of human blastoids from primed human pluripotent stem cells"

**The PDF file includes:**

Materials and Methods
Figs. S1 to S13
Tables S1 to S6
References 1 to 16

### Materials and methods

Culture medium

*Primed hPSC culture medium*

mTeSR-1 medium (Stem Cell Technologies, Vancouver, Canada) [1] is supplemented with 10 μM Y27632 (Wako, Osaka, Japan) [2].

*Naïve hPSC culture medium*

Naïve PXGL medium [3] was prepared as a 1:1 mixture of DMEM/F12 (Sigma-Aldrich, St. Louis, MO, USA) and neurobasal medium (Thermo Fisher Scientific, Waltham, MA, USA) supplemented with 2 mM L-glutamine (Thermo Fisher Scientific), 0.1 mM 2-mercaptoethanol (Wako), 0.5% (v/v) N2 supplement (Thermo Fisher Scientific), 1.0% (v/v) B27 supplement (ThermoFisher Scientific), 0.5% (v/v) penicillin/streptomycin (Wako), 10 ng/mL human leukemia inhibitory factor (hLIF, Nacalai Tesque, Inc, Kyoto, Japan), 250 μM L-ascorbic acid (Sigma), 10 μg/mL recombinant human insulin (Sigma), 1 μM PD0325901 (Wako), 2 μM XAV939 (Selleck Biotech, Tokyo, Japan), 2 μM Gö6983 (Cayman Chemical, Michigan, USA), and 10 μM Y27632.

*TSC medium*

TSC medium [4] contains DMEM/F12 with GlutaMAX (ThermoFisher Scientific) supplemented with 0.3% (w/v) bovine serum albumin (BSA) (Sigma-Aldrich), 0.2% (v/v) fetal bovine serum (FBS; Cell Culture Bioscience, Tokyo, Japan), and 0.5% (v/v) penicillin/streptomycin, 1% ITS-X supplement (ThermoFisher Scientific), 0.1 mM 2-mercaptoethanol, 1.5 μg/mL L-ascorbic acid, 5 μM Y27632, 2 μM CHIR99021 (Stemgent, Cambridge, MA, USA), 0.5 μM A83-01 (Wako), 1 μM SB431542(Wako), 50 ng/mL human recombinant EGF (PeproTech, Rocky Hill, NJ), and 0.8 mM valproic acid (VPA; Cayman Chemical).

*IVC1 medium*

IVC1 medium contains DMEM/F12, supplemented with 20% (v/v) FBS, 1% ITS-X supplement, 2 mM L-glutamine, 0.5% (v/v) penicillin/streptomycin, 25 μM *N*-acetyl-l-cysteine (Cayman Chemical), 8 nM β-estradiol (Cayman Chemical), and 200 ng/mL progesterone (Nacalai Tesque).

*IVC2 medium*

IVC2 medium contains DMEM/F12, supplemented with 30% knockout serum replacement (KSR; ThermoFisher Scientific), 1% ITS-X supplement, 2 mM L-glutamine, 0.5% (v/v) penicillin/streptomycin, 25 μM *N*-acetyl-l-cysteine, 8 nM β-estradiol, and 200 ng/mL progesterone.

*iBlastoid medium*

iBlastoid medium [5] contain 2:1:1 mixture of IVC1 medium, iBlastoid basal medium 1 (50:50 mixture of DMEM/F-12 and neurobasal medium, supplemented with 2 mM L-glutamine, 0.1 mM 2-mercaptoethanol, 0.5% N2 supplement, 1% B27 supplement, and 1% penicillin/streptomycin) and iBlastoid basal medium 2 (DMEM/F-12, GlutaMAX supplemented with 0.2% FBS, 0.1 mM 2-mercaptoethanol, 1% ITS-X supplement, 1.5 μg/mL l-ascorbic acid, 0.5% penicillin/streptomycin, and 0.3% BSA,) supplemented with 2 μM

CHIR99021, 0.5 μM A83-01, 0.8 mM VPA, 50 ng/mL EGF 1 μM SB431542, and 10 ng/mL BMP4 (R&D Systems, Minneapolis, MN, USA).

hESC culture

Human ESCs were used according to the guidelines provided by the ethical committee of Kyoto University (#ES3-9 and ES3-29). The K1-OCT4-eGFP cell line was obtained from Eihachiro Kawase [6]. Before culturing, 1.3 % (v/v) hESC-certified Matrigel (Corning, NY, USA) in DMEM/F12 was coated onto a culture dish and incubated 24 hours at 4 °C. Excess Matrigel was removed, and the coated dish was washed with fresh DMEM/F12. From here, the mTeSR-1 medium was used for daily culturing of hPSCs. For passaging, cells were dissociated with TrypLE Express (Life Technologies, Carlsbad, CA, USA) for 3 min at 37 °C, and then harvested through a cell strainer to remove undesired aggregates from the cell suspension. The cells were centrifuged at 200 × $g$ for 3 min, resuspended in the mTeSR-1 medium, and counted using a NucleoCounter NC-200 (Chemetec, Baton Rouge, LA, USA). To prevent apoptosis of dissociated hPSCs on day 1, mTeSR-1 medium supplemented with 10 μM Y27632 was used. After day 1, mTeSR-1 medium without Y27632 was used with daily medium changes. The cells were maintained in an incubator at 37 °C with 5 % (v/v) $CO_2$.

Atomic force microscope (AFM) nano-indentation for determination of gel elasticity

Indentation experiments were performed using an AFM (NanoWizard3, JPK Instruments, Berlin, Germany) mounted on a microscope (Olympus IX71, Tokyo, Japan). Cantilevers with a nominal spring constant $k = 0.1$ N/m were used, where the tips were modified with a borosilicate colloidal probe with a radius of $R = 5$ μm (CP-qp-CONT-BSG, sQUBE, Bickenbach a.d.B, Germany). PNIPAAm-PEG (HG) gels were kept at 37 °C in DMEM and indented after the system reached thermal equilibrium. For each sample, three independent areas of 100 × 100 μm$^2$ were selected, and 5 × 5 positions in each area indented at a tip indentation speed of 1 μm/s. The force curves obtained were analyzed by a self-written algorithm using Igor Pro (WaveMetrics, Portland, OR, USA). The elastic modulus was determined from the force-indentation ($F$-$\delta$) relationship using the modified Hertz model for sphere–plane contact [7,8].

$$F = \frac{4}{3}\frac{E}{1-\mu^2}\sqrt{R\delta^3}$$

where $E$ is the bulk elastic modulus (Young's modulus) and the Poisson's ratio, $\mu = 0.5$, is set to a constant value. The contact point between the cantilever tip and the sample was determined during the fitting process according to a previous report [9].

Generation of human blastoids in hydrogel (HG)

Human ESCs cultured on Matrigel-coated dishes in mTeSR-1 medium were harvested with TrypLE Express (Thermo Fisher Scientific) for 3 min at 37 °C and transferred to a 15 mL tube. The dissociated hPSCs were resuspended in mTeSR-1 medium supplemented with 10 μM of Y27632 and transferred to AggreWell 400 plates (Stem Cell Technologies) at 1.5 × 10$^5$ cells per well. After culturing at 37 ° C with 5 % (v/v) $CO_2$ for 24 hours, the hPSC aggregates were resuspended in the HG solution (DMEM with 10 % [w/v] PNIPAAm-β-PEG hydrogel [HG; Mebiol Inc., Hiratsuka, Japan]) at 4 °C. Next, 260 μL of an hPSC aggregate suspension with HG was transferred to the wells of a 24-well plate, and DMEM supplemented with 10 % (v/v) FBS was added at 37 °C. After 3 days, the iBlastoid medium was changed daily, and the cells maintained at 37 °C with 5 % (v/v) $CO_2$.

Quantitative RT-PCR

The primers used in this study are listed in **Table S3**. Total RNA was purified from cells using the RNeasy Mini Kit (Qiagen, Hilden, Germany), and 2.5 µg reverse transcribed into complementary DNA (cDNA) using the PrimeScript RT kit (TaKaRa Bio, Shiga, Japan). For the qRT-PCR, a reaction mixture of 25 µL containing 20 ng cDNA, 12.5 µL SYBR Premix Ex Taq II (Tli RNaseH Plus; TaKaRa Bio), 2 µL of 10 µM primers, and 0.5 µL ROX reference dye was introduced into each tube. The PCR conditions consisted of an initial denaturation at 95 °C for 30 s, followed by 40 cycles of 95 °C for 5 s and 60 °C for 31 s on an Applied Biosystems 7300 real-time PCR system (Applied Biosystems, Foster City, CA, USA). The DDCt method was used to calculate the relative mRNA expression levels of target genes. Housekeeping genes were used as loading control.

Immunocytochemistry

Cells were fixed with 4 % (v/v) paraformaldehyde in PBS for 60 min at 25 °C and then permeabilized with 25 mM glycine in PBS for 5 min, and finally, with 0.5 % (w/v) Triton X-100 (MP Biomedicals, CA, USA) in PBS for 1 hour at 25 °C. Next, the cells were blocked at 4 °C for 16 hours in PBS containing 5 % (v/v) normal donkey serum (Wako), 3 % (w/v) BSA, and 0.1 % (v/v) Triton X-100. After blocking, the cells were incubated with primary antibodies in a blocking buffer at 4 °C for 16 hours. Next, the cells were incubated with a secondary antibody and 300 nM 4′,6-diamidino-2-phenylindole (DAPI; Wako) in a blocking buffer at 25 °C for 2 hours. All the antibodies used in this study are listed in **Table S2**.

Image acquisition

Confocal microscopy was performed using an Andor Dragonfly 502 system (Andor, UK), built on a Nikon ECLIPSE Ti2-E inverted fluorescence microscope (Nikon, Tokyo, Japan). The system was equipped with a Zyla 4.2 Plus scientific complementary metal-oxide-semiconductor (sCMOS) camera (Andor) driven by Fusion (Andor), a CFI Plan apochromat lambda 10× dry/0.45 NA (Nikon), CFI Super Fluor 20× dry/0.75 NA (Nikon), motorized XY stage and stage piezo (Applied Scientific Instrumentation, MO, USA), and three lasers (405, 488, 561, and 637 nm).

Preparation of cell samples for single-cell RNA sequencing

For single-cell RNA sequencing (scRNA-seq), HG-blastoids in a dish were placed on ice and washed with 0.04 % BSA in PBS twice at 4 °C. Approximately 200 HG-blastoids were collected and examined to determine their blastocyst-like morphology (i.e., intact spherical shape, intact cell aggregates in a cyst, and blastocyst-like cavity). Collected HG-blastoids were trypsinized with Accumax (Nacalai Tesque) for 40 min at 37 °C and washed with 0.04 % BSA in PBS twice. As a control, hESCs cultured in a 35 mm dish were trypsinized with Accumax for 5 min at 37 °C. Dissociated cells were applied to a chromium controller (10× Genomics, Pleasanton, CA, USA) to encapsulate the cells, following the manufacturer's instructions (Chromium Next GEM Single Cell 3' Reagent Kits v3.1 [Dual Index]). Obtained RNA was quantified using qRT-PCR with the Collibri™ Library Quantification Kit (Thermo Fisher Scientific) according to the manufacturer's instructions. The PCR conditions consisted of an initial denaturation at 95 °C for 2 min, followed by 35 cycles of 95 °C for 30 s and 60 °C for 45 s on a StepOnePlus system (Applied Biosystems, Waltham, MA, USA). For each sample, 1.8 pM was applied using the NextSeq 500 Mid Output Kit v2.5 (Illumina, San Diego, CA, USA). Sequencing was carried out with Illumina Next Seq 500 (Illumina) using a pair-end (Read1:28 bp, Read2:90 cycles) sequencing strategy to obtain 20,000 reads per cell. Chromium barcodes were used for demultiplexing with the CellRanger program (10× Genomics; v6.1.1, https://support.10xgenomics.com/single-cell-gene-expression/software/overview/welcome). Alignment and UMI counting were performed using CellRanger to map 373,641,788 sequencing reads onto a custom version of the Ensembl GRCh38 reference genome.

Blastoid scRNA-seq

To investigate individual cell identities in HG-blastoids, scRNA-seq analyses were carried out using RStudio (v1.4.1717) and Seurat (v4.1.2) [10,11], and the data were visualized using ggplot2 (v3.3.5) [12]. The scRNA-seq results were loaded into the Seurat software, and the cells were filtered with unique feature counts over 5,000 or less than 200. Moreover, cells were filtered when the mitochondrial counts were over 5 %. After filtering, each sample contained at least 2,000 cells for further analyses. After normalizing the data in Seurat and conducting a principal component analysis (PCA), a UMAP was generated using five dimensions. Unsupervised clustering was then performed using the "FindClusters" function, with a resolution of 0.1. To find the differentially expressed features among the clusters, the "FindMarkers" function was performed with the minimum threshold of fold changes set to 0.25. CellMarker 2.0 (http://bio-bigdata.hrbmu.edu.cn/CellMarker/index.html) was also used to name the identified clusters. Trajectory analysis based on scRNA-seq data was performed in R using Monocle3 [16–18]. To transform Seurat object to Monocle3, we set cluster cells resolution of 1e-5. We selected the EPI1 cluster as the starting point for Monocle3.

Integration of human blastocyst, hESCs, and iBlastoids scRNA-seq data

To compare our scRNA-seq data with previously published scRNA-seq data, Seurat was used to integrate the data integrated using the "FindIntegrationAnchors" function, followed by "IntegrateData" function. Datasets included the following: Petropoulos et al. (ArrayExpress# E-MTAB-3929; https://www.ebi.ac.uk/arrayexpress/experiments/E-MTAB-3929/) [13], Liu et al (GEO#: GSE156596) [5], Yanagida et al. (GEO#: GSE171820) [14], and Yu et al. (GEO#: GSE150578) [15]. After normalizing the data in Seurat and conducting a PCA, a UMAP was generated using 20 dimensions. Unsupervised clustering was performed using the "FindClusters" function, with a resolution of 0.5. To find the differentially expressed features among clusters, the "FindMarkers" function was performed with the minimum threshold of fold changes set to 0.25.

Derivation of naive and primed bPSCs from HG-blastoids

For the derivation of naive bPSCs, blastoids were collected, dissociated with Accumax, and seeded onto mitomycin C-treated mouse embryonic fibroblasts (MEFs; Cosmo Bio, Sapporo, Japan) in iBlastoid medium supplemented with 10 µM Y27632. After culturing at 37 °C with 5 % (v/v) $CO_2$ for 24 hours, the medium was replaced with naive PXGL medium. To maintain naive PSCs, cells were passaged every 5–6 days at a 1:3 sub-culture ratio using TrypLE Express on MEFs, with daily medium changes.

For the derivation of primed bPSCs, blastoids were collected, dissociated with Accumax, and seeded onto Matrigel-coated dishes in iBlastoid medium supplemented with 10 µM Y27632. After 24 hours, the medium was replaced with mTeSR-1 medium. To maintain primed bPSCs, cells were passaged every 3–4 days at a 1:10–1:20 sub-culture ratio using TrypLE Express on Matrigel-coated dishes, with daily medium changes.

Derivation of trophoblast stem cells (bTSCs)

For the derivation of bTSCs, after collecting blastoids, dissociated cells were seeded on 5 µg/mL collagen IV (Sigma-Aldrich)-coated plates in iBlastoid medium supplemented with 10 µM Y27632. After 24 hours, the medium was replaced with the TSC medium supplemented with 5 µM Y27632. Cells were passaged every 4–6 days at 1:3–1:5 sub-culture ratios using TrypLE Express on 5 µg/mL collagen IV-coated plates with medium changes every other day.

*In vitro* implantation assay

The in *vitro* implantation assay was performed according to previously published protocols [14,16]. Briefly, blastocysts were collected on day 9 and sedimented for 5–10 min to remove the majority of cell debris within the supernatant. Blastoids were then transferred onto glass-bottom tissue culture plates (Iwaki, Japan) coated with Geltrex (Thermo Fisher Scientific), and cultured in IVC1 medium with daily medium replacement at 37 °C with 5 % (v/v) $CO_2$. On day 2, the culture medium was replaced with the IVC2 medium. HG-blastoids were cultured up to day 4.5 with daily medium replacement and collected for immunocytochemistry, qRT-PCR analysis, and hCG ELISA.

hCG ELISA

To confirm hCG production by both HG-blastoids and attached HG-blastoids, the medium was collected from both HG-blastoids (day 9) and attached HG-blastoids (day 9 + 4.5) and stored at -80 °C. hCG ELISA was performed using a human hCG ELISA kit (Abcam, Cambridge, UK) according to the manufacturer’s protocol.

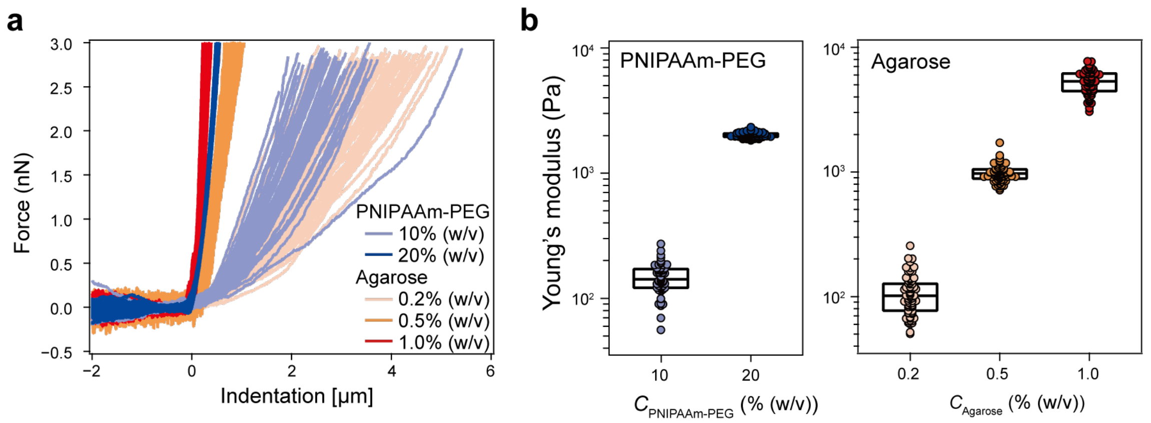

**Fig. S1| a,** Overlays of the force-indentation curves obtained from PNIPAAm-PEG gels (HG) at 10% (w/v) (light blue) and 20% (w/v) (dark blue), and agarose gels at 0.2% (w/v) (light orange), 0.5% (w/v) (orange), and 1.0% (w/v) (red). Indentation measurements were performed at 75 positions on each gel. All the curves that were fitted with the modified Hertz model were overlaid. **b,** Distribution of Young's modulus *E* corresponding to PNIPAAm-PEG gels at 10 and 20% (w/v), and agarose gels at 0.2, 0.5, and 1.0% (w/v). The middle line, bottom, and top of each box plot correspond to the median, first, and third quartiles, respectively.

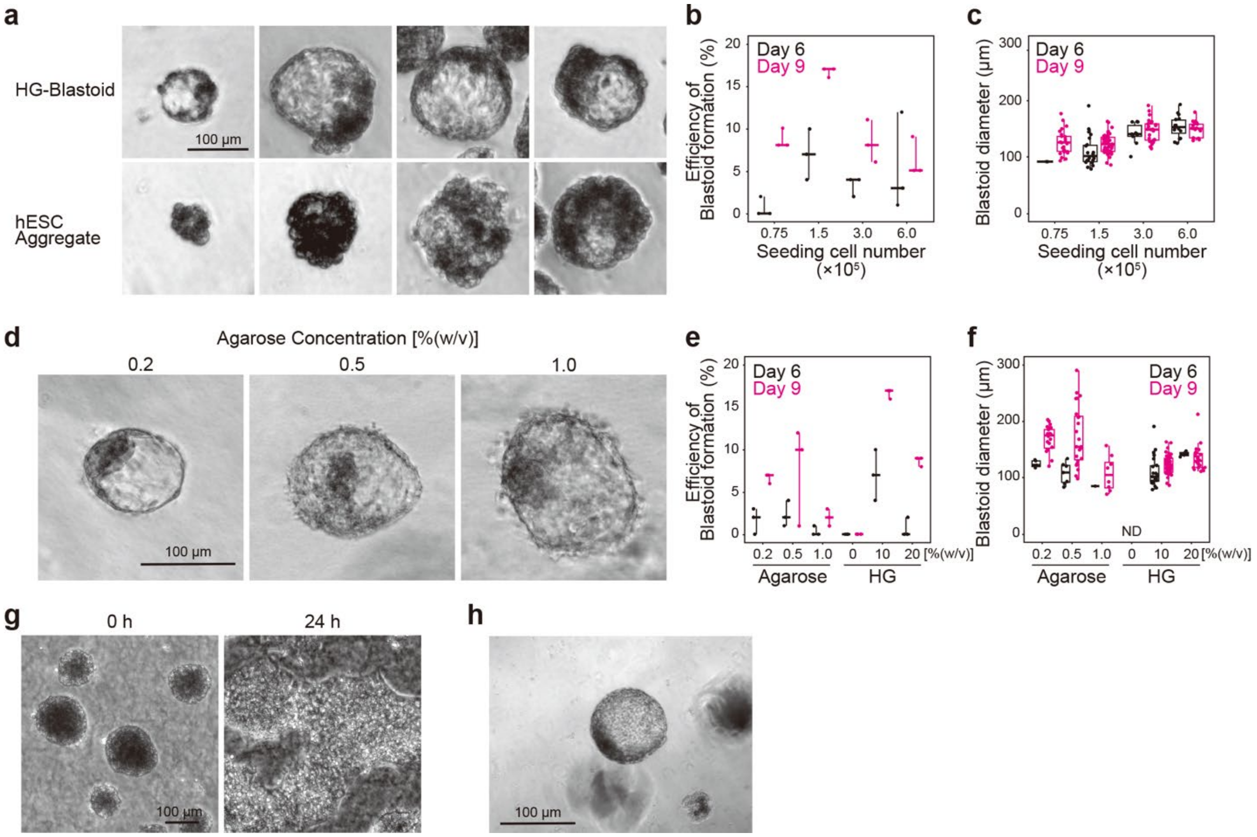

**Fig. S2| a,** Micrographs of HG-blastoids and hESC aggregates of K1-OCT4-eGFP cells generated in HG at 10% (w/v). **b,** Efficiencies to generate HG-blastoids with initial seeding cell numbers at 0.75, 1.5, 3.0, and 6.0 × $10^5$ cells in a well of the AggreWell 400 plate at the end of HG-blastoid generation process shown in **Fig. 2A**. The lines show the minimum, median and maximum values. **C,** Size distribution of HG-blastoids with initial seeding cell numbers at 0.75, 1.5, 3.0, and 6.0 × $10^5$ cells in a well of the AggreWell 400 plate at the end of HG-blastoid generation process shown in **Fig. 2A**. Center lines show the medians; box limits indicate the 25$^{th}$ and 75$^{th}$ percentiles; whiskers extend 1.5 times the interquartile range from the 25$^{th}$ and 75$^{th}$ percentiles. **d,** Migrographs of HG-blastoids generated in agarose gel at 0.2, 0.5. and 1.0 % (w/v). **e,** Generation efficiency of HG-blastoids in 0.2, 0.5, and 1.0 % (w/v) agarose gel and 0, 10, and 20% (w/v) HG. The lines show the minimum, median and maximum values. **f,** Size distribution of HG-blastoids in 0, 10, and 20% (w/v) HG in 0.2, 0.5, and 1.0 % (w/v) agarose gel and at Day 9 of blastoid generation. **g,** Micrographs of hPSC aggregates cultured in collagen gel at 2 mg/mL. **h,** A micrograph of HG-blastoids derived from 585A1 hiPSCs generated in HG at 10% (w/v) at Day 9.

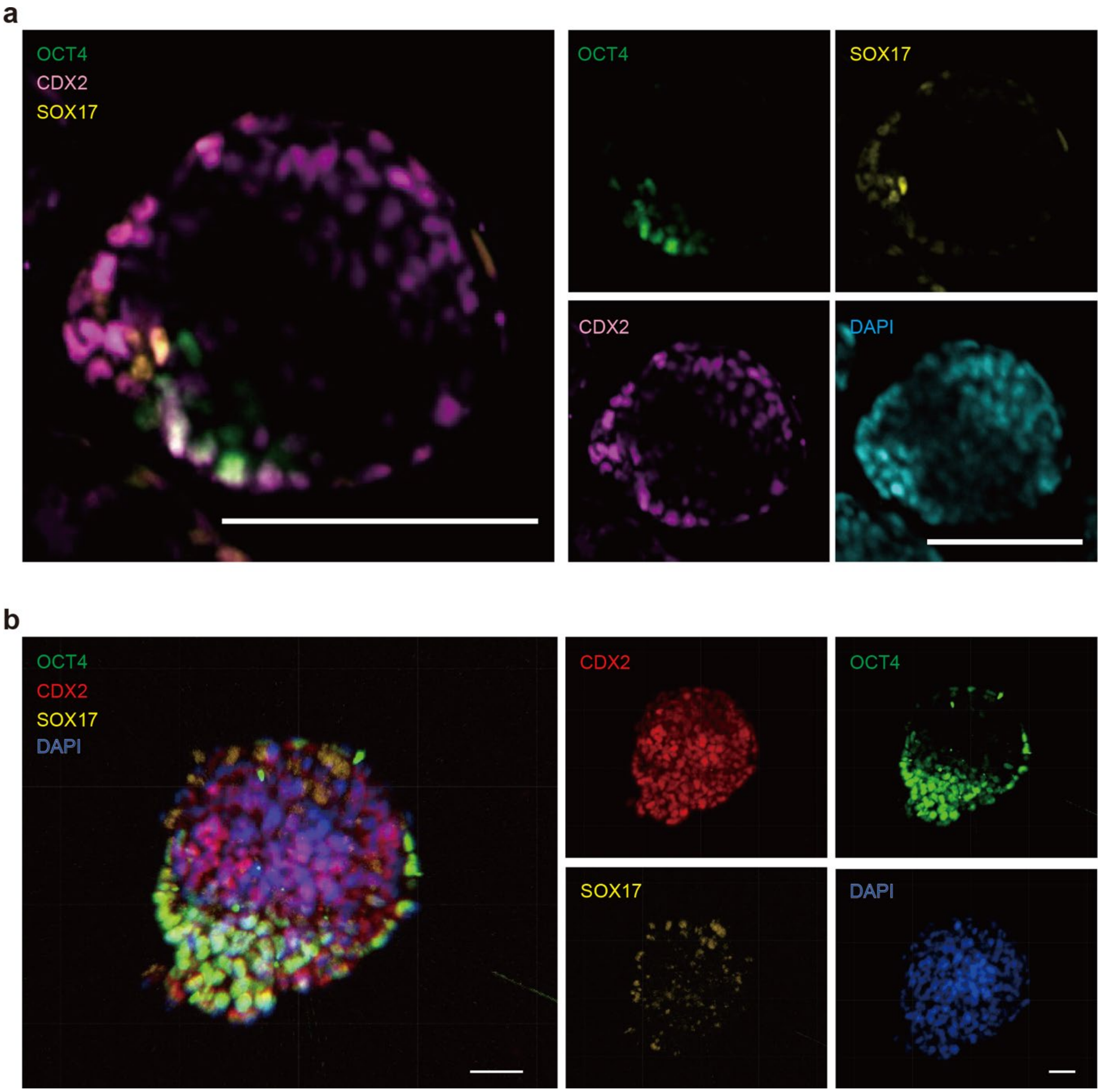

**Fig. S3| a,** Micrographs of whole-mount immunocytochemistry to observe the typical markers of epiblast (OCT4), trophectoderm (CDX2) and primitive endoderm (SOX17) in HG-Blastoids of K1-OCT4-eGFP cells. DAPI was used for nuclear staining, as shown in **Fig. 1D**. All scale bars represent 100 μm. **b,** Micrographs of whole-mount immunocytochemistry for OCT4, CDX2 and SOX17 in HG-Blastoids of 585A1 hiPSCs. DAPI was used for nuclear staining. All scale bars represent 20 μm.

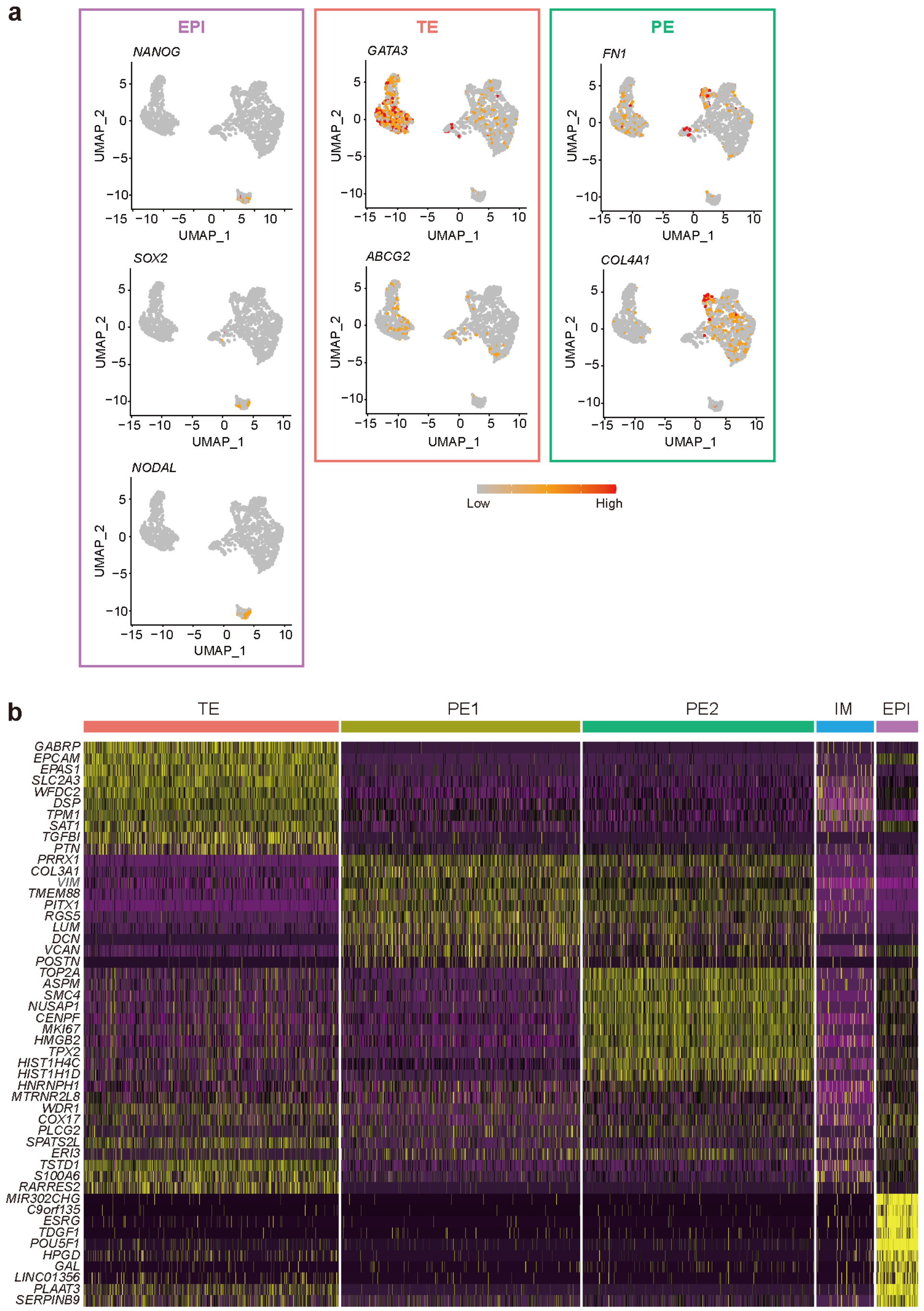

**Fig. S4| a,** UMAP for 2072 K1-OCT4-eGFP cells in HG-Blastoid sc-RNA-seq library to show the expression of marker genes of EPI, TE, and PE. **b,** Differentially expressed genes in EPI, TE, PE, and IM cells.

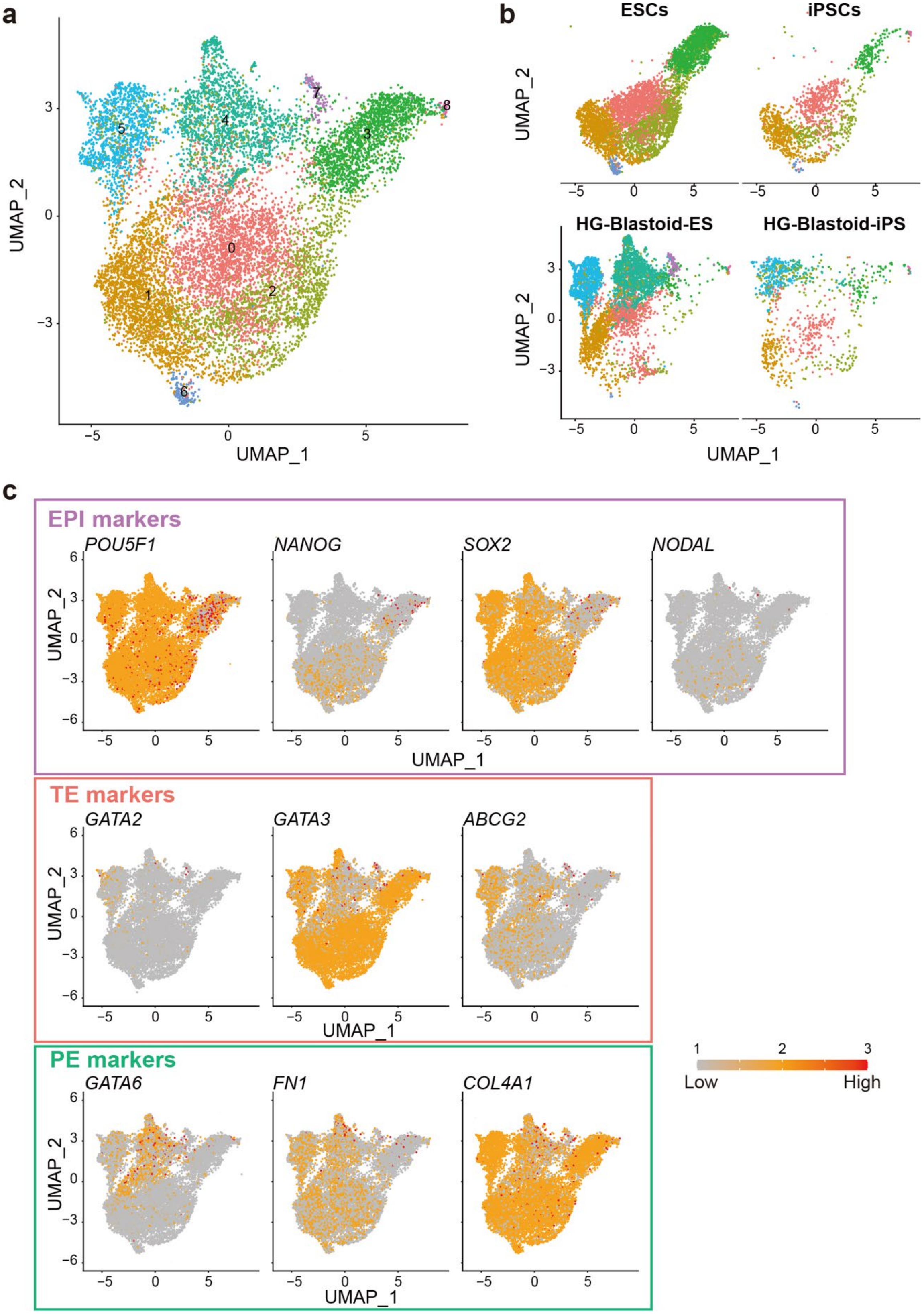
a
b
ESCs
iPSCs
HG-Blastoid-ES
HG-Blastoid-iPS
UMAP_1
UMAP_2
c
EPI markers
POU5F1
NANOG
SOX2
NODAL
TE markers
GATA2
GATA3
ABCG2
PE markers
GATA6
FN1
COL4A1
Low
High

**Fig. S5| Comparison of single-cell transcriptomic patterns of hESCs and HG-Blastoids. a,** Integrated UMAP of scRNA-seq data sets of of K1-OCT4-eGFP hESCs (ESCs), HG-Blastoids from K1-OCT4-eGFP cells (HG-Blastoid-ES), 585A1 hiPSCs (iPSCs) and HG-Blastoids from 585A1 cells (HG-Blastoid-iPS). UMAP showed 9 clusters. **b,** Split UMAP for ESCs, iPSCs, HG-Blastoid-ES and HG-Blastoid-iPS. Interestingly, both hESCs iPSCs showed four major cell populations (Clusters #0, #1, #2, and #3). Both HG-Blastoid-ES and HG-Blastoid-iPS have large overlapped cellular populations, but HG-Blastoid-ES had a large cellular population in Cluster #4, but not HG-Blastoid-iPS. Moreover, HG-Blastoid-ES and HG-Blastoid-iPS lost cell populations in Clusters #2 and #3, compared with ESCs and iPSCs, while Cluster #5 was appeared in HG-Blastoid-ES and HG-Blastoid-iPS. **c,** UMAP feature plots to show the expression of marker genes associated with EPI (*POU5F1, NANOG*, *SOX2*, and *NODAL*), TE (*GATA2, GATA3*, and *ABCG2*) and PE (*GATA6*, *FN1*, and *COL4A1*). Cluster #4 showed the gene expression associated with PE markers, and Cluster #5 showed TE-like transcriptomic signatures. These clusters were not observed in K1-OCT4-eGFP hESCs nor 585A1 hiPSCs and thus, appeared via HG-assisted blastoid generation.

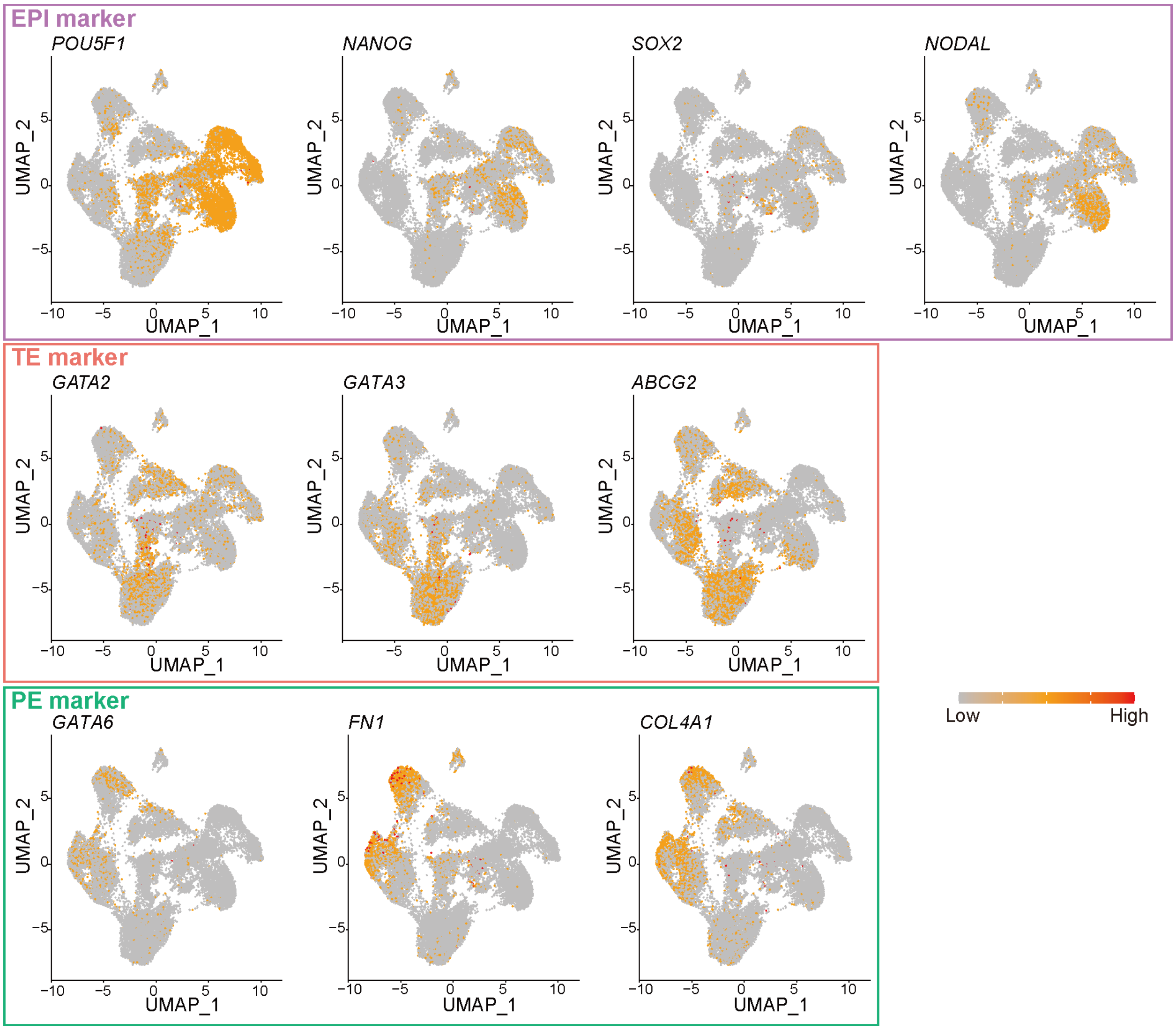

**Fig. S6 |** UMAP feature plot of the expression of marker genes of EPI (*POU5F1*, *NANOG*, *SOX2*, and *NODAL*), TE (*GATA2, GATA3*, and *ABCG2*) and PE (*GATA6*, *FN1*, and *COL4A1*) of the integrated scRNA-seq data sets integrated with HG-Blastoids derived from K1-OCT4-eGFP cells, human E5-E7 blastocysts [13], and published blastoids (Yu Nature 2021, Liu Nature 2021, Yanagida Cell Stem Cell 2021), shown in **Fig. 2D and E**.

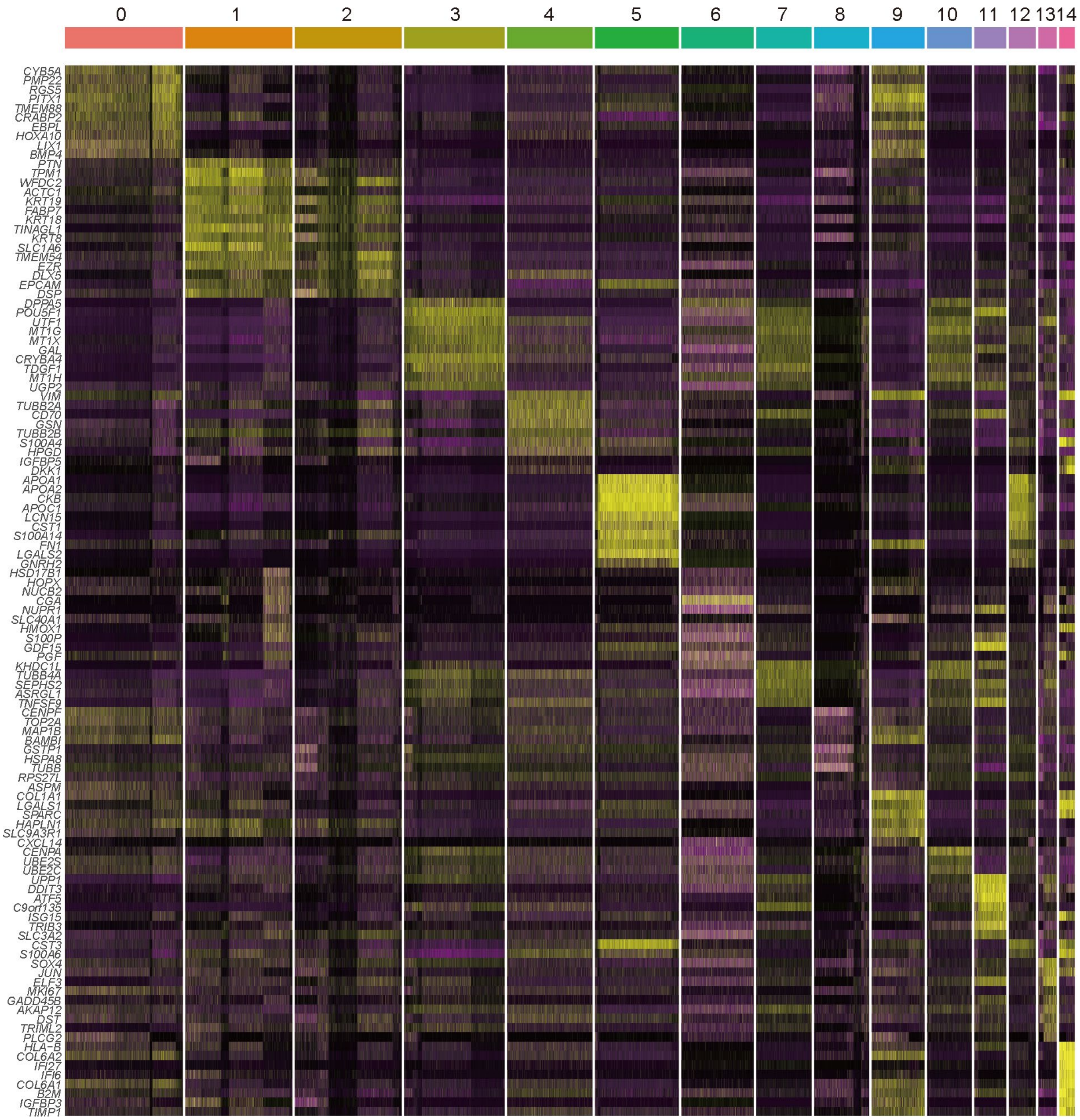

**Fig. S7 |** Heatmap of top 10 activated genes in each cluster compared to the other cells in UMAP of integrated sc-RNA-seq data sets shown in **Fig. 2D and E**.

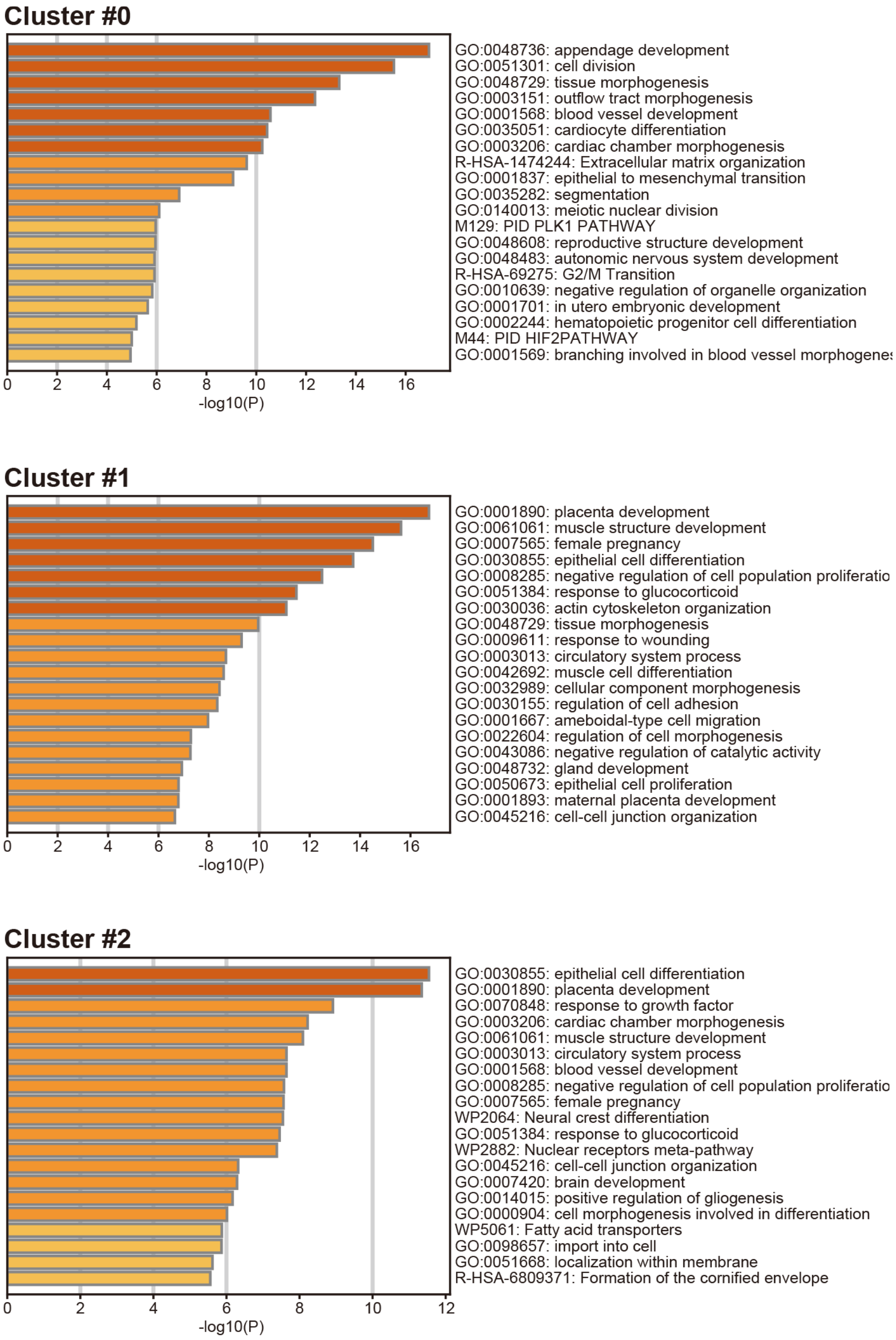

**Fig. S8|** Enriched gene ontology terms in each cluster compared to the other cells in UMAP of integrated sc-RNA-seq data sets shown in **Fig. 2D and E**.

### Cluster #3

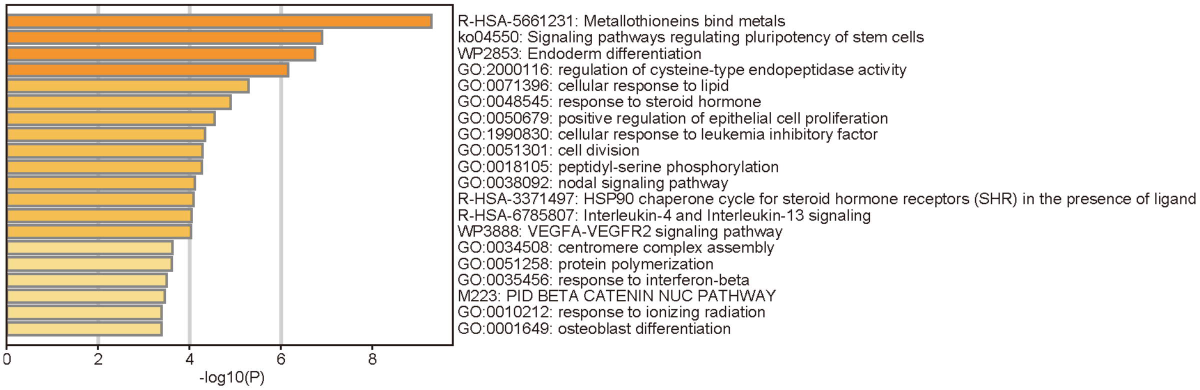

### Cluster #4

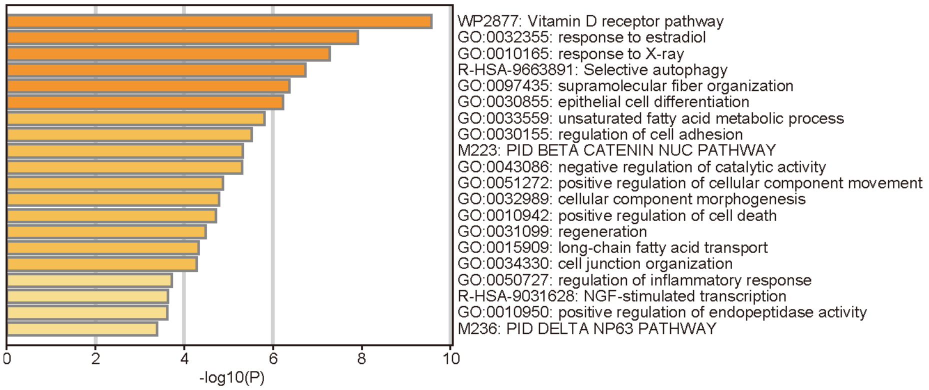

### Cluster #5

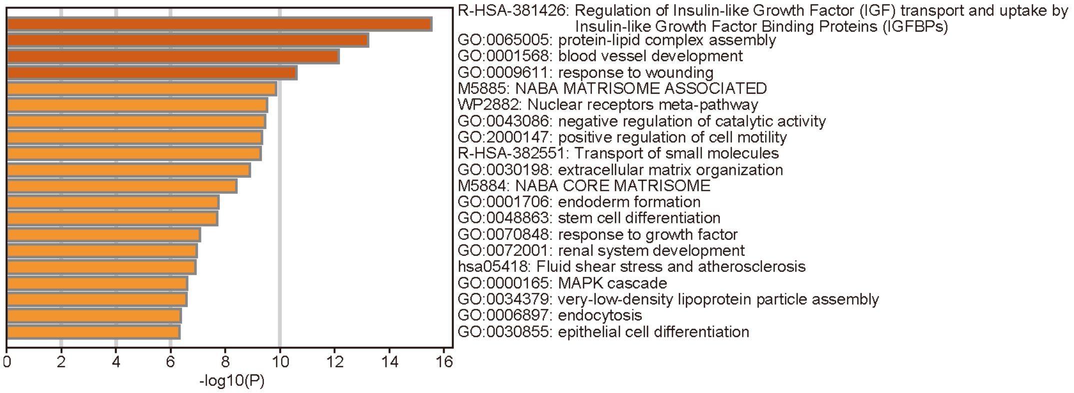

**Fig. S8 | (cont.)**

### Cluster #6

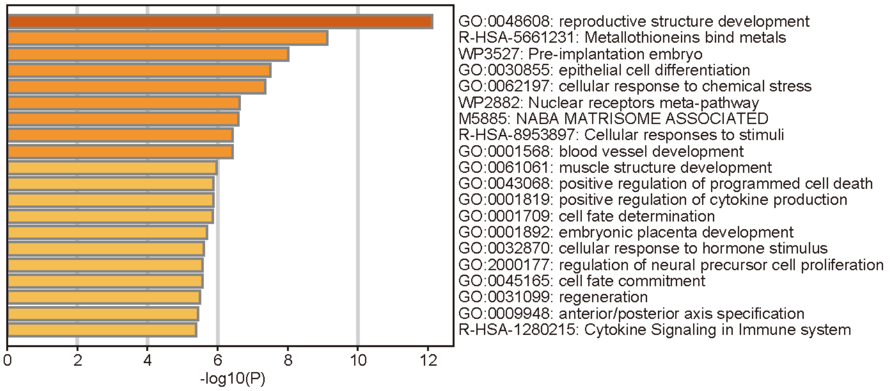

### Cluster #7

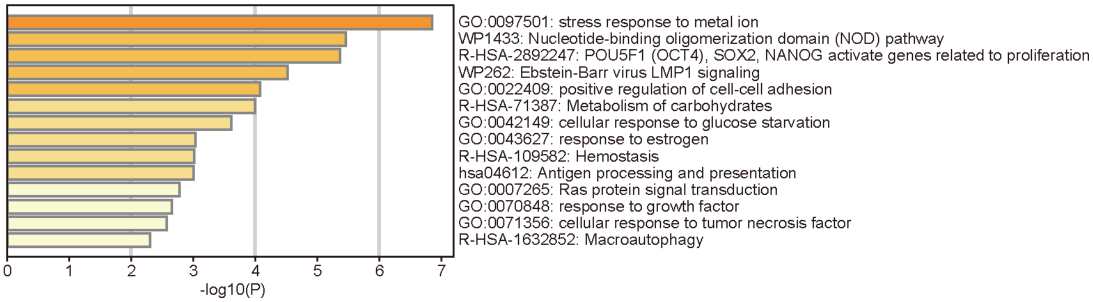

### Cluster #8

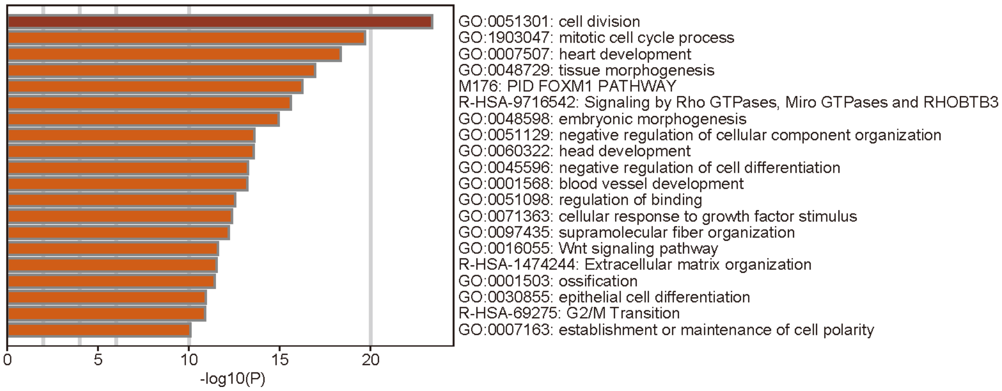

**Fig. S8 | (cont.)**

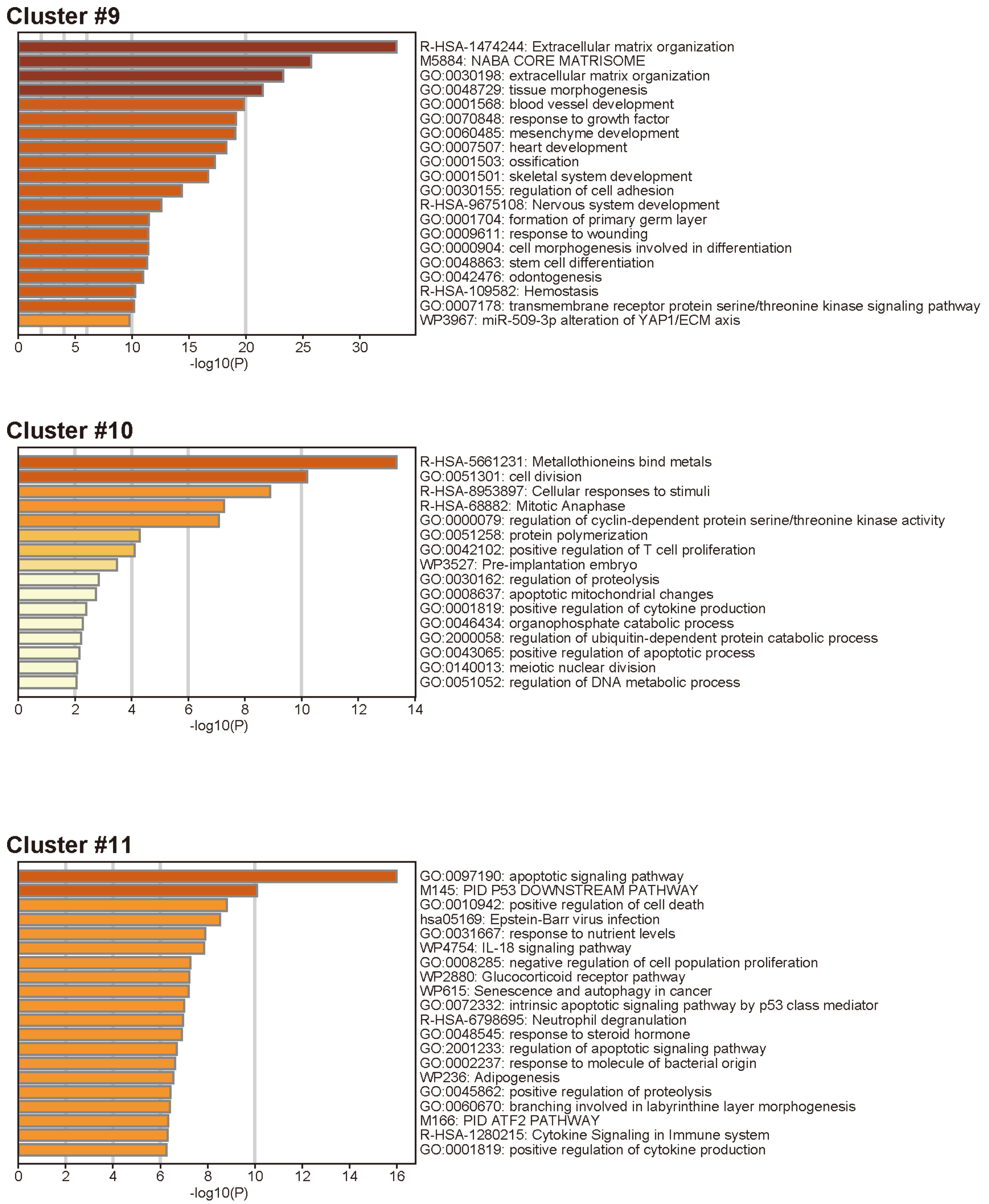
Cluster #9
R-HSA-1474244: Extracellular matrix organization
M5884: NABA CORE MATRISOME
GO:0030198: extracellular matrix organization
GO:0048729: tissue morphogenesis
GO:0001568: blood vessel development
GO:0070848: response to growth factor
GO:0060485: mesenchyme development
GO:0007507: heart development
GO:0001503: ossification
GO:0001501: skeletal system development
GO:0030155: regulation of cell adhesion
R-HSA-9675108: Nervous system development
GO:0001704: formation of primary germ layer
GO:0009611: response to wounding
GO:0000904: cell morphogenesis involved in differentiation
GO:0048863: stem cell differentiation
GO:0042476: odontogenesis
R-HSA-109582: Hemostasis
GO:0007178: transmembrane receptor protein serine/threonine kinase signaling pathway
WP3967: miR-509-3p alteration of YAP1/ECM axis
0
5
10
15
20
25
30
-log10(P)
Cluster #10
R-HSA-5661231: Metallothioneins bind metals
GO:0051301: cell division
R-HSA-8953897: Cellular responses to stimuli
R-HSA-68882: Mitotic Anaphase
GO:0000079: regulation of cyclin-dependent protein serine/threonine kinase activity
GO:0051258: protein polymerization
GO:0042102: positive regulation of T cell proliferation
WP3527: Pre-implantation embryo
GO:0030162: regulation of proteolysis
GO:0008637: apoptotic mitochondrial changes
GO:0001819: positive regulation of cytokine production
GO:0046434: organophosphate catabolic process
GO:2000058: regulation of ubiquitin-dependent protein catabolic process
GO:0043065: positive regulation of apoptotic process
GO:0140013: meiotic nuclear division
GO:0051052: regulation of DNA metabolic process
0
2
4
6
8
10
12
14
-log10(P)
Cluster #11
GO:0097190: apoptotic signaling pathway
M145: PID P53 DOWNSTREAM PATHWAY
GO:0010942: positive regulation of cell death
hsa05169: Epstein-Barr virus infection
GO:0031667: response to nutrient levels
WP4754: IL-18 signaling pathway
GO:0008285: negative regulation of cell population proliferation
WP2880: Glucocorticoid receptor pathway
WP615: Senescence and autophagy in cancer
GO:0072332: intrinsic apoptotic signaling pathway by p53 class mediator
R-HSA-6798695: Neutrophil degranulation
GO:0048545: response to steroid hormone
GO:2001233: regulation of apoptotic signaling pathway
GO:0002237: response to molecule of bacterial origin
WP236: Adipogenesis
GO:0045862: positive regulation of proteolysis
GO:0060670: branching involved in labyrinthine layer morphogenesis
M166: PID ATF2 PATHWAY
R-HSA-1280215: Cytokine Signaling in Immune system
GO:0001819: positive regulation of cytokine production
0
2
4
6
8
10
12
14
16
-log10(P)

**Fig. S8 | (cont.)**

### Cluster #12

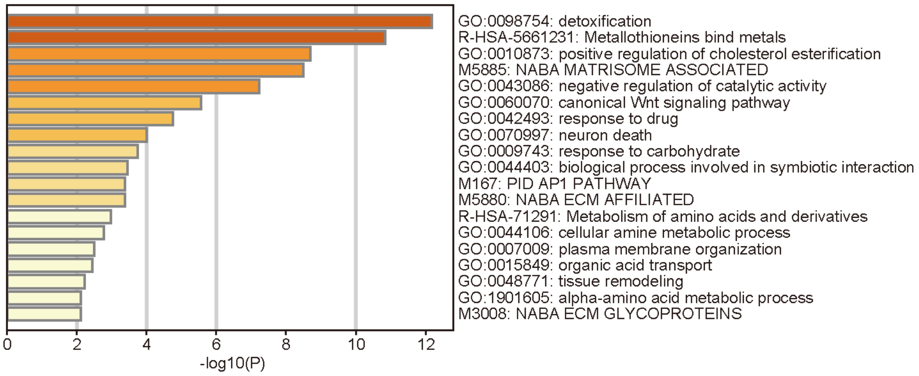

### Cluster #13

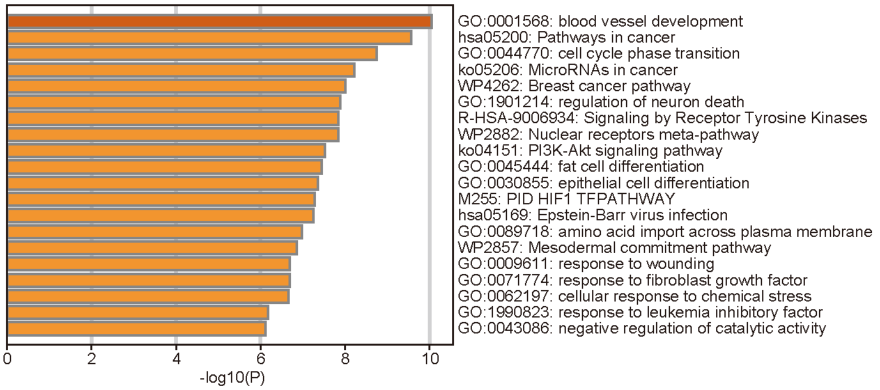

### Cluster #14

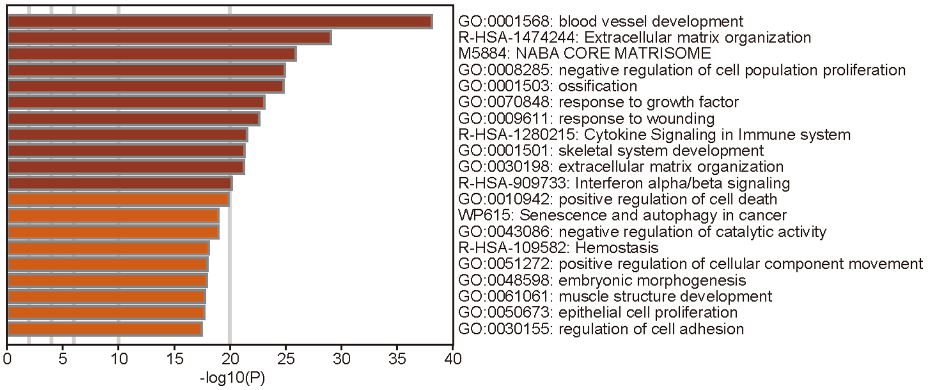

**Fig. S8 | (cont.)**

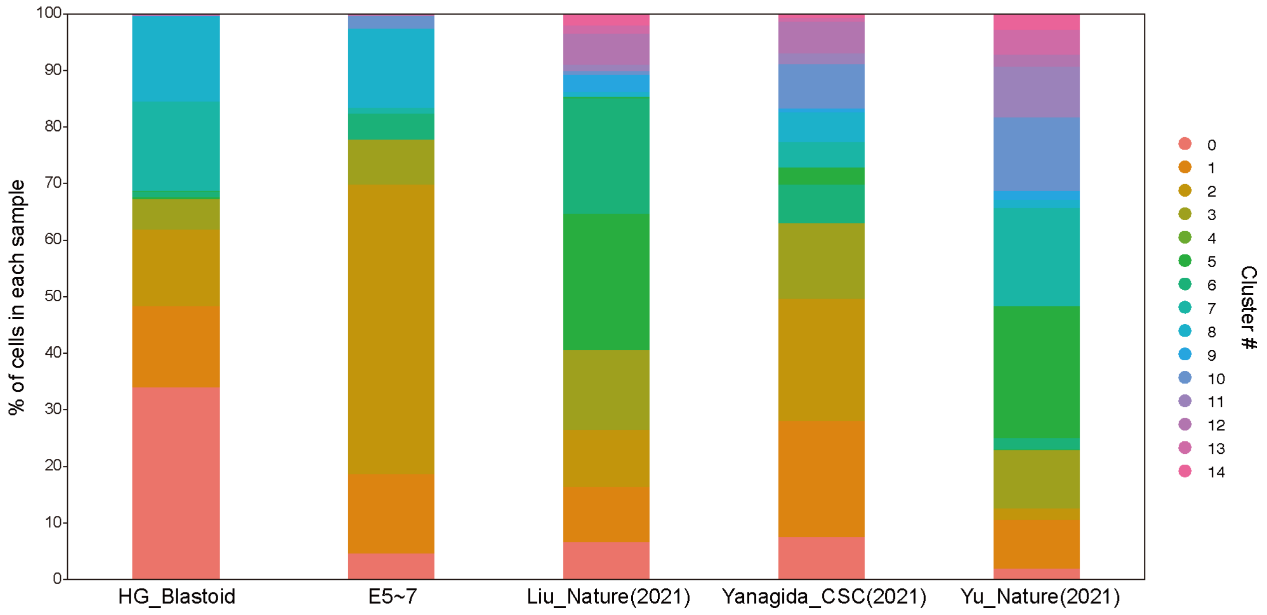

**Fig. S9 |** A bar graph of percentiles of cells in each sample defined in UMAP shown in **Fig. 2D and E**.

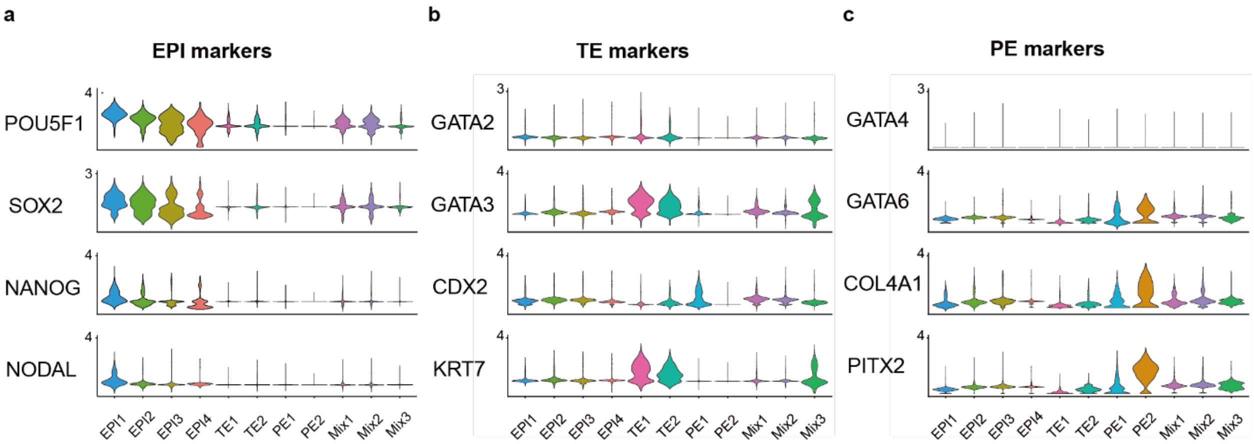

**Fig. S10 | Expression levels of blastoid markers (EPI, TE, and PE) in each cluster. a**, Expression levels of EPI markers (*POF5F1*, *SOX2*, *NANOG*, and *NODAL*) in each cluster. **b,** Expression levels of TE markers (*GATA2*, *GATA3*, *CDX2*, and *KRT7*) in each cluster. **c,** Expression levels of PE markers (*GATA4*, *GATA6*, *COL4A1*, and *PITX2*) in each cluster.

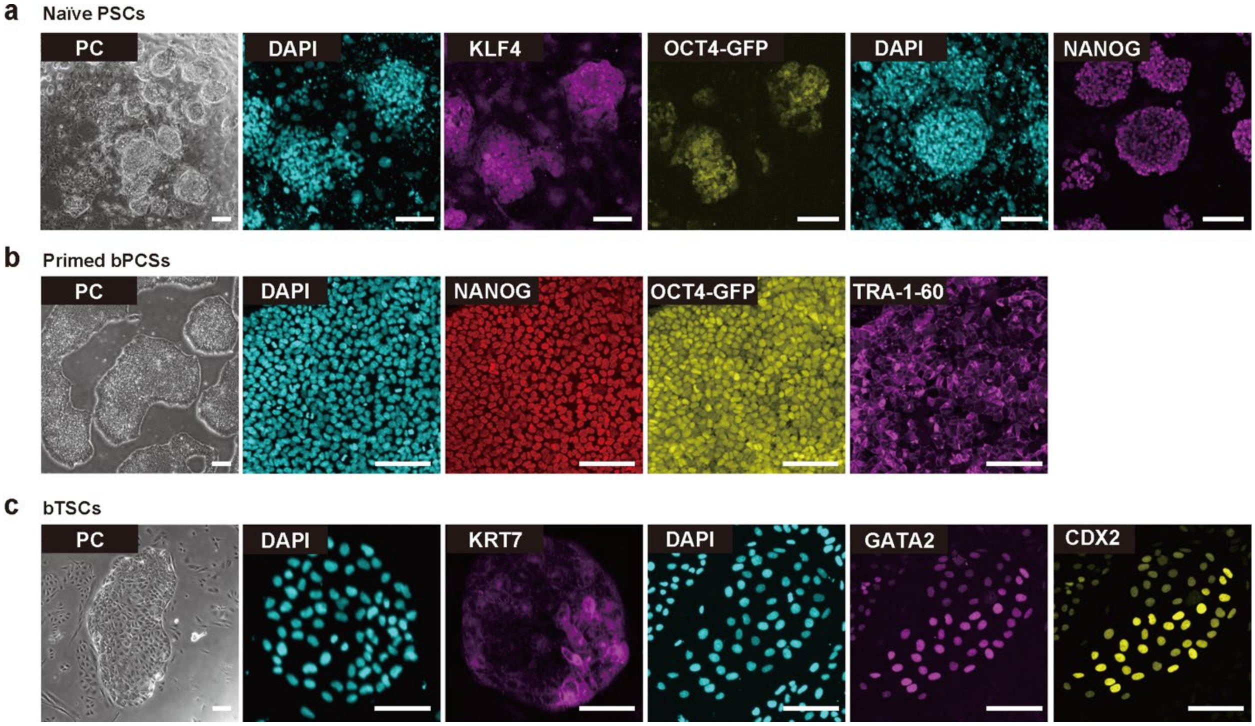

**Fig. S11 | Derivation of stem cells from HG-Blastoids derived from of K1-OCT4-eGFP hESCs. a,** Phase-contrast and fluorescent micrographs of immunocytochemistry of naive bPSC stained with KLF4, NANOG, and DAPI. GFP was expressed under the regulation of *OCT4* promoter. **b,** Phase-contrast and fluorescent micrographs of immunocytochemistry of primed bPSCs stained with NANOG, TRA1-60, and DAPI. GFP was expressed under the regulation of *OCT4* promoter. **c,** Phase-contrast and fluorescent micrographs of immunocytochemistry of bTSCs stained with KRT7, GATA2, and DAPI. Scale bar = 100 µm.

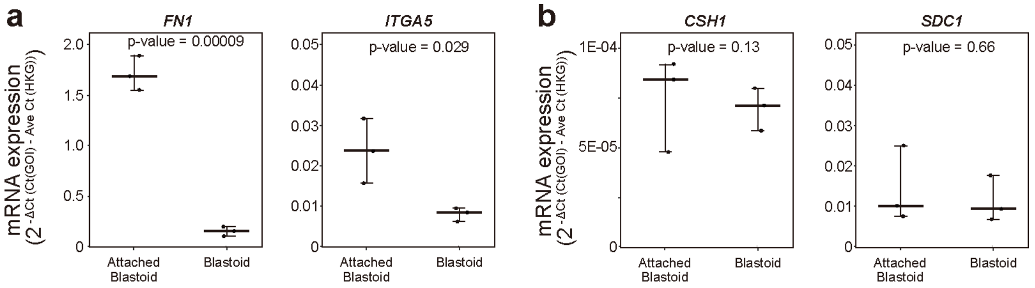

**Fig. S12 |** Quantitative RT-PCR analysis for the expression of genes associated with markers of **a**, extravillous cytotrophoblasts (EVTs; *FN1* and *ITGA5*) and **b**, syncytiotrophoblasts (STs; *CSH1* and *SDC1*). n = 3.

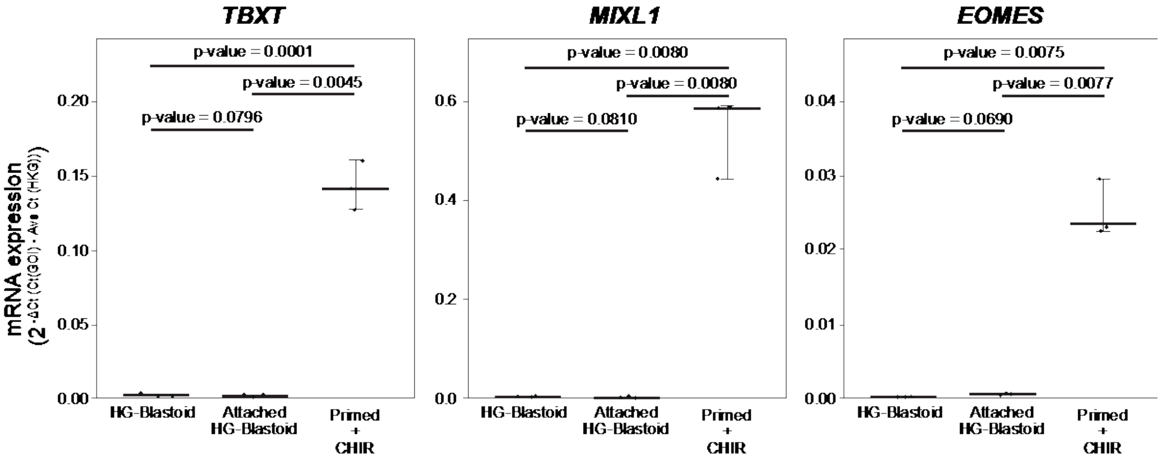

**Fig. S13 |** Quantitative RT-PCR analysis for the expression of genes associated with primitive streak markers, *TBXT*, *MIXL1*, and *EOMES*. Boxplot in which the median of each group is indicated by a black line (25$^{th}$–75$^{th}$ interquartile range). n = 3.

**Table S1 | Young's modulus *E* of agarose and mebiol gels.**

| Gel | Concentration (% (w/v)) | Young's modulus *E* (kPa) |
|---|---|---|
| Agarose | 0.2 | 0.11 ± 0.04 |
| | 0.5 | 0.98 ± 0.16 |
| | 1.0 | 5.35 ± 1.11 |
| Mebiol | 10 | 0.15 ± 0.04 |
| | 20 | 2.01 ± 0.10 |

### Table S2 | Statistical data of generation efficiency and size distribution of HG-blastoid of primed K1-OCT4-eGFP hESCs.

**Seeding cell number**

**Generation efficiency of HG-blastoids**

**D6 Blastoid**

| Group 1 | Group 2 | p-value |
|---|---|---|
| 7.5×10^4 | 1.5×10^5 | 0.180 |
| 7.5×10^4 | 3.0×10^5 | 0.773 |
| 7.5×10^4 | 6.0×10^5 | 0.390 |
| 1.5×10^5 | 3.0×10^5 | 0.574 |
| 1.5×10^5 | 6.0×10^5 | 0.929 |
| 3.0×10^5 | 6.0×10^5 | 0.886 |

**D9 Blastoid**

| Group 1 | Group 2 | p-value |
|---|---|---|
| 7.5×10^4 | 1.5×10^5 | 0.003 |
| 7.5×10^4 | 3.0×10^5 | 0.996 |
| 7.5×10^4 | 6.0×10^5 | 0.447 |
| 1.5×10^5 | 3.0×10^5 | 0.002 |
| 1.5×10^5 | 6.0×10^5 | 0.001 |
| 3.0×10^5 | 6.0×10^5 | 0.565 |

**D6 Blastoid vs D9 Blastoid**

| Group 1 (D6) | Group 2 (D9) | p-value |
|---|---|---|
| 7.5×10^4 | 7.5×10^4 | 0.001 |
| 1.5×10^5 | 1.5×10^5 | 0.005 |
| 3.0×10^5 | 3.0×10^5 | 0.035 |
| 6.0×10^5 | 6.0×10^5 | 0.797 |

**Size distribution of HG-blastoids**

**D6 Blastoid**

| Group 1 | Group 2 | p-value |
|---|---|---|
| 7.5×10^4 | 1.5×10^5 | 0.686 |
| 7.5×10^4 | 3.0×10^5 | 0.041 |
| 7.5×10^4 | 6.0×10^5 | 0.005 |
| 1.5×10^5 | 3.0×10^5 | 0.007 |
| 1.5×10^5 | 6.0×10^5 | 0.000 |
| 3.0×10^5 | 6.0×10^5 | 0.546 |

**D9 Blastoid s**

| Group 1 | Group 2 | p-value |
|---|---|---|
| 7.5×10^4 | 1.5×10^5 | 0.914 |
| 7.5×10^4 | 3.0×10^5 | 0.001 |
| 7.5×10^4 | 6.0×10^5 | 0.000 |
| 1.5×10^5 | 3.0×10^5 | 0.000 |
| 1.5×10^5 | 6.0×10^5 | 0.000 |
| 3.0×10^5 | 6.0×10^5 | 0.965 |

**D6 Blastoid vs D9 Blastoid**

| Group 1 (D6) | Group 2 (D9) | p-value |
|---|---|---|
| 7.5×10^4 | 7.5×10^4 | 0.027 |
| 1.5×10^5 | 1.5×10^5 | 0.013 |
| 3.0×10^5 | 3.0×10^5 | 0.413 |
| 6.0×10^5 | 6.0×10^5 | 0.518 |

**Agarose gel and hydrogel**

**Generation efficiency of HG-blastoids**

**D6 Blastoid**

| Group 1 | Group 2 | p-value |
|---|---|---|
| 0.2% Agarose gel | 0.5% Agarose gel | 0.998 |
| 0.2% Agarose gel | 1.0% Agarose gel | 0.856 |
| 0.2% Agarose gel | 0% Mebiol gel | 0.632 |
| 0.2% Agarose gel | 10% Mebiol gel | 0.363 |
| 0.2% Agarose gel | 20% Mebiol gel | 0.935 |
| 0.5% Agarose gel | 1.0% Agarose gel | 0.467 |
| 0.5% Agarose gel | 0% Mebiol gel | 0.294 |
| 0.5% Agarose gel | 10% Mebiol gel | 0.484 |
| 0.5% Agarose gel | 20% Mebiol gel | 0.758 |
| 1.0% Agarose gel | 0% Mebiol gel | 0.918 |
| 1.0% Agarose gel | 10% Mebiol gel | 0.346 |
| 1.0% Agarose gel | 20% Mebiol gel | 1.000 |
| 0% Mebiol gel | 10% Mebiol gel | 0.294 |
| 0% Mebiol gel | 20% Mebiol gel | 0.918 |
| 10% Mebiol gel | 20% Mebiol gel | 0.346 |

**D9 Blastoid**

| Group 1 | Group 2 | p-value |
|---|---|---|
| 0.2% Agarose gel | 0.5% Agarose gel | 0.995 |
| 0.2% Agarose gel | 1.0% Agarose gel | 0.257 |
| 0.2% Agarose gel | 0% Mebiol gel | 0.054 |
| 0.2% Agarose gel | 10% Mebiol gel | 0.003 |
| 0.2% Agarose gel | 20% Mebiol gel | 0.911 |
| 0.5% Agarose gel | 1.0% Agarose gel | 0.121 |
| 0.5% Agarose gel | 0% Mebiol gel | 0.023 |
| 0.5% Agarose gel | 10% Mebiol gel | 0.008 |
| 0.5% Agarose gel | 20% Mebiol gel | 0.995 |
| 1.0% Agarose gel | 0% Mebiol gel | 0.911 |
| 1.0% Agarose gel | 10% Mebiol gel | 0.000 |
| 1.0% Agarose gel | 20% Mebiol gel | 0.054 |
| 0% Mebiol gel | 10% Mebiol gel | 0.000 |
| 0% Mebiol gel | 20% Mebiol gel | 0.010 |
| 10% Mebiol gel | 20% Mebiol gel | 0.018 |

**D6 Blastoid vs D9 Blastoid**

| Group 1 (D6) | Group 2 (D9) | p-value |
|---|---|---|
| 0.2% Agarose gel | 0.2% Agarose gel | 0.006 |
| 0.5% Agarose gel | 0.5% Agarose gel | 0.202 |
| 1.0% Agarose gel | 1.0% Agarose gel | 0.067 |
| 0% Mebiol gel | 0% Mebiol gel | NA |
| 10% Mebiol gel | 10% Mebiol gel | 0.005 |
| 20% Mebiol gel | 20% Mebiol gel | 0.072 |

**Size distribution of HG-blastoids**

D6 Blastoid

| Group 1 | Group 2 | p-value |
|---|---|---|
| 0.2% Agarose gel | 0.5% Agarose gel | 0.881 |
| 0.2% Agarose gel | 1.0% Agarose gel | 0.586 |
| 0.2% Agarose gel | 10% Mebiol gel | 0.460 |
| 0.2% Agarose gel | 20% Mebiol gel | 0.298 |
| 0.5% Agarose gel | 1.0% Agarose gel | 0.586 |
| 0.5% Agarose gel | 10% Mebiol gel | 1.000 |
| 0.5% Agarose gel | 20% Mebiol gel | 0.298 |
| 1.0% Agarose gel | 10% Mebiol gel | 0.762 |
| 1.0% Agarose gel | 20% Mebiol gel | 0.737 |
| 10% Mebiol gel | 20% Mebiol gel | 0.474 |

**D9 Blastoid**

| Group 1 | Group 2 | p-value |
|---|---|---|
| 0.2% Agarose gel | 0.5% Agarose gel | 1.000 |
| 0.2% Agarose gel | 1.0% Agarose gel | 0.003 |
| 0.2% Agarose gel | 10% Mebiol gel | 0.000 |
| 0.2% Agarose gel | 20% Mebiol gel | 0.000 |
| 0.5% Agarose gel | 1.0% Agarose gel | 0.028 |
| 0.5% Agarose gel | 10% Mebiol gel | 0.000 |
| 0.5% Agarose gel | 20% Mebiol gel | 0.034 |
| 1.0% Agarose gel | 10% Mebiol gel | 0.658 |
| 1.0% Agarose gel | 20% Mebiol gel | 0.292 |
| 10% Mebiol gel | 20% Mebiol gel | 0.201 |

**D6 Blastoid vs D9 Blastoid**

| Group 1 (D6) | Group 2 (D9) | p-value |
|---|---|---|
| 0.2% Agarose gel | 0.2% Agarose gel | 0.000 |
| 0.5% Agarose gel | 0.5% Agarose gel | 0.013 |
| 1.0% Agarose gel | 1.0% Agarose gel | NA |
| 10% Mebiol gel | 10% Mebiol gel | 0.063 |
| 20% Mebiol gel | 20% Mebiol gel | 0.339 |

### Table S3 | Antibody list

| Antibody | Cat. # | Clone | RRID | Supplier | Working conc. (µg/mL) | Working dilution | Applications |
|---|---|---|---|---|---|---|---|
| OCT4 | sc-5279 | C-10 | AB_628051 | Santa Cruz | 2 | 1/100 | IHC |
| NANOG | 4903S | D73G4 | AB_10559205 | Cell Signaling | 0.1 | 1/100 | IHC |
| SOX2 | NB110- 37235 | - | AB_792070 | Novus Biologicals | 20 | 1/50 | IHC |
| CDX2 | ab76541 | EPR2764 Y | AB_1523334 | Abcam | 1.6 | 1/500 | IHC |
| GATA2 | WH0002624M1 | 2D11 | AB_1841726 | Merck | 10 | 1/100 | IHC |
| GATA6 | MAB1700-SP | 222228 | AB_2108899 | R&D systems | 5 | 1/200 | IHC |
| SOX17 | AF1924 | - | AB_355060 | R&D systems | 10 | 1/40 | IHC |
| KLF4 | 11880-1-AP | - | AB_10640807 | Proteintech | 7 | 1/100 | IHC |
| KLF17 | HPA024629 | - | AB_1668927 | Atlas Antibodies | 1 | 1/100 | IHC |
| TRA1-60 | sc-21705 | TRA-1-60 | AB_628385 | Santa Cruz | 2 | 1/100 | IHC |
| KRT7 | HPA007272 | - | AB_1079181 | Atlas Antibodies | 1 | 1/200 | IHC |
| Phalloidin (F-actin)-iFluor 594 Reagent | ab176757 | - | - | Abcam | - | 1/1000 | IHC |
| Podocalyxin | MAB1658 | # 222328 | AB_2165985 | R&D Systems | 10 | 1/60 | IHC |
| aPKC | sc-17781 | H-1 | AB_628148 | Santa Cruz | 0.4 | 1/500 | IHC |
| MMP-2 | #40994 | D4M2N | AB_2799191 | Cell Signaling | 3.5 | 1/200 | IHC |
| hCGb | ab9582 | 5H4-E2 | AB_296507 | Abcam | 10 | 1/100 | IHC |
| HLA-G | ab7759 | MEM-G/1 | AB_306053 | Abcam | 10 | 1/100 | IHC |
| Alexa Fluor 488 Donkey anti-goat IgG | 705-546-147 | | AB_2340430 | Jackson ImmunoResearch | | 1/1000 | IHC |
| Alexa Fluor 488 Donkey anti-rabbit IgG | 711-546-152 | | AB_2340619 | Jackson ImmunoResearch | | 1/1000 | IHC |
| Alexa Fluor 594 Donkey anti-rabbit IgG | 711-586-152 | | AB_2340622 | Jackson ImmunoResearch | | 1/1000 | IHC |
| Alexa Fluor 647 Donkey anti-rabbit IgG | 711-606-152 | | AB_2340625 | Jackson ImmunoResearch | | 1/1000 | IHC |
| Alexa Fluor 594 Donkey anti-mouse IgG | 715-586-150 | | AB_2340857 | Jackson ImmunoResearch | | 1/1000 | IHC |
| Alexa Fluor 647 Donkey anti-mouse IgG | 715-606-150 | | AB_2340865 | Jackson ImmunoResearch | | 1/1000 | IHC |
| Alexa Fluor 488 Donkey anti-mouse IgG | 715-546-150 | | AB_2340849 | Jackson ImmunoResearch | | 1/1000 | IHC |
| Rhodamine (TRITC)-Donkey-anti-goat IgG(H+L) | 705-025-147 | | AB_2340389 | Jackson ImmunoResearch | | 1/1000 | IHC |
| AlexaFlour647 Donkey anti-Mouse IgM | 715-606-020 | | AB_2340864 | Jackson ImmunoResearch | | 1/1000 | IHC |
| Alexa Fluor 488 Donkey anti-goat IgG | 705-546-147 | | AB_2340430 | Jackson ImmunoResearch | | 1/1000 | IHC |
| Alexa Fluor 488 Donkey anti-rabbit IgG | 711-546-152 | | AB_2340619 | Jackson ImmunoResearch | | 1/1000 | IHC |
| Alexa Fluor 594 Donkey anti-rabbit IgG | 711-586-152 | | AB_2340622 | Jackson ImmunoResearch | | 1/1000 | IHC |
| Alexa Fluor 647 Donkey anti-rabbit IgG | 711-606-152 | | AB_2340625 | Jackson ImmunoResearch | | 1/1000 | IHC |

**Table S4 | Primer list**

| Primers | Sequence 5' → 3' |
|---|---|
| CSH1_Forward | CATGACTCCCAGACCTCCTTCT |
| CSH1_Reverse | ATTTCTGTTGCGTTTCCTCCAT |
| SDC1_Forward | CTGCCGCAAATTGTGGCTAC |
| SDC1_Reverse | TGAGCCGGAGAAGTTGTCAGA |
| CGB_Forward | ACCGTCAACACCACCATCTGTG |
| CGB_Reverse | GAAGCGCACATCGCGGTAGTTG |
| ITGA1_Forward | GCTCCTCACTGTTGTTCTACG |
| ITGA1_Reverse | CGGGCCGCTGAAAGTCATT |
| FN1_Forward | CGGTGGCTGTCAGTCAAAG |
| FN1_Reverse | AAACCTCGGCTTCCTCCATAA |
| ITGA5_Forward | GGCTTCAACTTAGACGCGGAG |
| ITGA5_Reverse | TGGCTGGTATTAGCCTTGGGT |
| TBXT_Forward | TATGAGCCTCGAATCCACATAGT |
| TBXT_Reverse | CCTCGTTCTGATAAGCAGTCAC |
| EOMES_Forward | GTGCCCACGTCTACCTGTG |
| EOMES_Reverse | CCTGCCCTGTTTCGTAATGAT |
| MIXL1_Forward | GGCGTCAGAGTGGGAAATCC |
| MIXL1_Reverse | GGCAGGCAGTTCACATCTACC |
| GAPDH_Forward | GCACCGTCAAGGCTGAGAAC |
| GAPDH_Reverse | TGGTGAAGACGCCAGTGGA |

**Table S5 | List of high expressed cell markers among clusters.**

See attached Table S5.

**Table S6 | List of high expressed cell markers and candidate cells in Mix3.**

See attached Table S6.
